# The Arabidopsis U1 snRNP regulates mRNA 3’-end processing

**DOI:** 10.1101/2023.09.19.558503

**Authors:** Anchilie F. Mangilet, Joachim Weber, Sandra Schüler, Irina Droste-Borel, Samuel Streicher, Thomas Schmutzer, Gregor Rot, Boris Macek, Sascha Laubinger

**Author notes:** Max Planck Institute for Plant Breeding Research (MPIPZ), 50829 Cologne Germany (Anchilie F. Mangilet).

## Abstract

The removal of introns by the spliceosome is a key gene regulatory mechanism in eukaryotes, with the U1 snRNP subunit of the spliceosome playing a crucial role in the early stages of splicing. Studies in metazoans show that the U1 snRNP also conducts splicing-independent functions, but the lack of genetic tools and knowledge about U1 snRNP-associated proteins have limited the study of such splicing-independent functions in plants. Here, we describe an RNA-centric approach that identified more than 200 proteins associated with the Arabidopsis U1 snRNP, among them mRNA cleavage and polyadenylation factors. The loss of U1 core components is linked to premature cleavage and polyadenylation within gene bodies and alternative polyadenylation site selection in 3’-UTRs. Overall, our work provides a comprehensive view of U1 snRNP interactors and reveals novel functions in regulating mRNA 3’-end processing in Arabidopsis, thus establishing the groundwork for a better understanding of non-canonical functions of plant U1 snRNPs.

## MAIN

In eukaryotes, the spliceosome removes intronic sequences in mRNAs and subsequently ligates exons to generate a functional mRNA. Five uridine-rich small nuclear ribonucleoprotein complexes (U1, U2, U4, U5, and U6 snRNPs) build the spliceosome ^1^. Each of the snRNPs is composed of a specific small nuclear RNA (snRNA) and protein subunits that are essential for the recognition of splicing signals embedded in the gene sequences ^2^. During the splicing process, the snRNPs assemble in a step-by-step and accurate manner. The recognition of the 5’ splice site (5’SS) by the U1 snRNP initiates the splicing process. Cryo-electron microscopy has facilitated a more detailed dissection of the U1 snRNP function, particularly in the early steps of the splicing reaction ^3–5^. The core U1 snRNP consists of a 165 nucleotide snRNA that forms four stem-loops, an Sm core ring (Sm-E, G, D3, B, D1, D2, and F), and three U1 core proteins (U1-A, U1-70K, and U1-C) ^6,7^. Accessory proteins specifically interact with the U1 core snRNP and aid splicing of weak 5’ splice sites^8–11^. In Arabidopsis, core and accessory proteins are conserved, and analyses of mutants lacking U1 accessory components such as LUC7, PRP39, PRP40, or PRP45 exhibit developmental defects ^12–19^. Surprisingly, while a flower-specific RNAi-knockdown of U1-70K shows developmental defects, two reports describing mutants for the U1 core components, U1-A and U-70K, did not find any drastic effects, ^20–22^. This is in stark contrast to the fact that U1 core components are essential genes in metazoans^23,24^ and it shows that several aspects of the function of the Arabidopsis U1 snRNP function in plants are not understood and remain to be answered.

The U1 snRNP is also particularly interesting because it is more abundant than other snRNPs and has early been thought to fulfill additional functions aside from splicing ^25^. Indeed, the U1 snRNP affects mRNA length through regulation of 3’-end processing, it controls promoter directionality, enhances transcription, and increases the speed of RNA polymerase II (RNAPII) and it is responsible for retaining lncRNA in the nucleus ^6,26–30^. Probably the best described function of the U1 snRNP is telescripting, by which the U1 snRNP prevents premature cleavage and polyadenylation in introns and thereby ensures transcription of full-length RNAs ^31^. Telescripting function is particularly important for long genes, which contain long introns and require intact U1 snRNP to complete transcription at canonical PAS ^32^. Environmental cues can also modulate telescripting activity and several human diseases can be linked to telescripting ^33–35^. Whether telescripting exists in plants, particularly in plants with rather small introns such as Arabidopsis, is currently not known. Mechanistically, the metazoans U1 snRNP forms a complex with cleavage and polyadenylation factors (CPAFs), U1-CPAF, which is distinct from U1 snRNP spliceosomal complexes ^36^. The U1-CPAF complex binds nascent RNAs in introns that contain U1 and CPAF binding sites, but the presence of the U1 snRNP in this complex blocks cleavage-stimulatory factors from joining the complex ^36^.

While numerous exciting non-canonical functions of metazoan snRNPs have been constantly discovered, comprehensive knowledge about the interactors of plant U1 snRNPs, as well as genetic tools to study the function of U1 snRNP in plants, is still lacking. In this study, we present the Arabidopsis U1 snRNP interactome and, in addition, generate genetic resources to investigate the non-canonical functions of the Arabidopsis U1 snRNP. Our findings demonstrate that the Arabidopsis U1 snRNP plays a splicing-independent role in 3’-end processing, as it features a telescripting function similar to metazoans, while also contributing to alternative polyadenylation in 3’UTRs, possibly coupled with a general function in RNAPII termination.

## RESULTS

### A compendium of Arabidopsis U1 snRNP-associated proteins

Despite the importance of the U1 snRNP in splicing and beyond, very little is known about the composition of the U1 snRNP or associated proteins and complexes in plants. To identify the proteins associated with a plant U1 snRNP complex, we applied “comprehensive identification of RNA-binding proteins by mass-spectrometry” (CHIRP-MS), which has been successfully applied to isolate proteins associated with the U1 snRNA or other non-coding RNAs^37^. We used a biotinylated U1 snRNA antisense probe to purify the Arabidopsis U1 snRNP and analyzed the purified sample by mass spectrometry (U1-IP-MS, Figure 1A). A short-distance cross-linking agent, formaldehyde, preserved also transient interactions of the U1 snRNP with other proteins and complexes during the purification procedure. To test whether we can indeed observe also dynamic interaction with this approach, we performed a similar experiment with an antisense oligonucleotide specific for the U2 snRNA (U2-IP-MS). The U1 snRNP, as part of the commitment complex, recruits the U2 snRNP for the formation of the A complex. Hence, we would expect a partially overlapping set of proteins associated with the U1 and the U2 snRNA. As a negative control, we performed an IP-MS experiment using an antisense *lacZ* oligonucleotide, the sequence of which is not expected to bind any RNA encoded in the Arabidopsis genome. Three biological replicates were prepared for each IP-MS experiment. In total, we were able to identify 908 proteins by MS (Figure 1B, complete lists on Supplemental Data Set 1).

**Figure 1:**
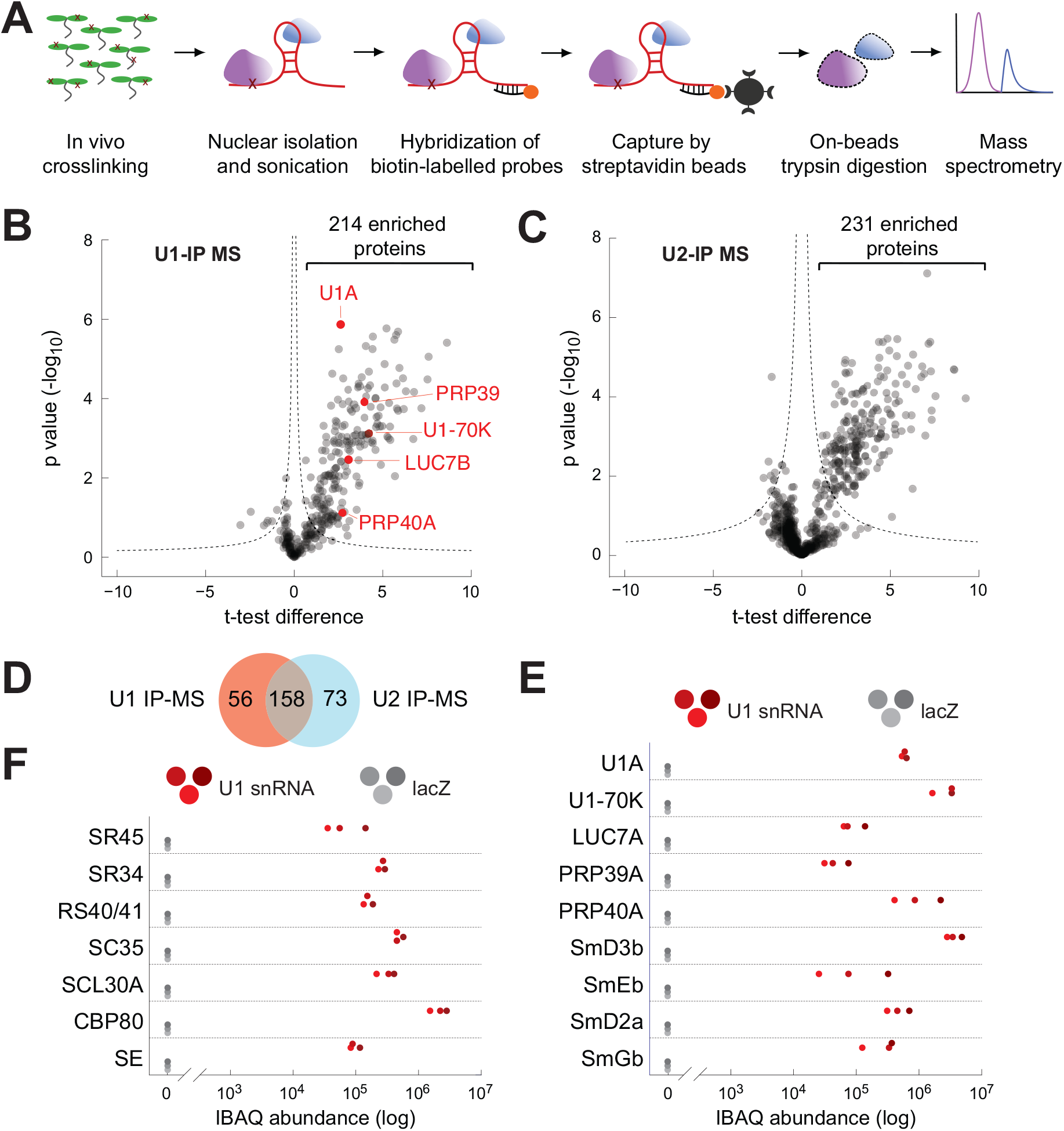
Identification of Arabidopsis U1 snRNP-associated proteins by U1-IP-MS. **A:** Schematic representation of the U1 snRNA immunoprecipitation followed by mass spectrometry (U1-IP-MS) experiment. **B, C:** Analysis of U1 snRNA (B) and U2 snRNA (C) associated proteins identified by IP-MS. Volcano plot of three biological replicates showing significantly enriched proteins immunoprecipitated with a U1 (B) or U2 (C) antisense oligonucleotide compared to a *lacZ* oligonucleotide (p-value < 0.01). Known U1-specific proteins are highlighted in red (B). **D:** Venn diagram depicting the overlap between significantly enriched proteins in U1-IP-MS and U2-IP-MS experiments. **E, F:** Abundance of specific proteins in U1-IP-MS experiments. The three red and grey dots represent intensity-based absolute quantification (IBAQ) values of three biological replicates using the U1 or the lacZ antisense oligonucleotide, respectively. Proteins known to be part of the U1 snRNP (E) and selected proteins that function in splicing and RNA processing (F) are shown.

We found 214 proteins significantly enriched in IPs with the U1 snRNA antisense probe (Figure 1B, Supplemental Table S1). With the U2 snRNA antisense probe, we retrieved 231 significantly enriched proteins (Figure 1C, Supplemental Table S2). 157 proteins were found to be associated with both the U1 and U2 snRNA antisense probe, while 56 and 73 proteins were specifically associated with the U1 and U2 snRNA antisense probe, respectively (Figure 1D). The large number of proteins that co-purified with the U1 and U2 snRNA antisense probes indicates that our approach was able to capture transient interactions that occur e.g. during the formation of the A complex. The effectiveness of the U1 snRNA IP is further supported by the successful enrichment of known U1 snRNP core and accessory components; we found known U1 snRNPs components such as U1-A, U1-70K, LUC7A/B, PRP39, PRP40A and Sm core proteins (SmB, SmD1, SmE, SmG) (Figure 1B, E, Supplemental Table S1). Not a single peptide of the above-mentioned protein was retrieved in the control IP experiments using the lacZ antisense probe (Figure 1E). U1-IP-MS also enriched splicing factors, many of which are known to interact with the U1 snRNP including the Serine/Arginine-rich (SR) proteins SR45, SR34, RS40/41, SC35, and SCL30A (Figure 1F, Supplemental Table S1). We also retrieved other splicing-related proteins, such as SERRATE and the nuclear cap-binding complex (nCBC), as well as components of the MOS4-associated complex (MAC), which is the homologue of the metazoan Nineteen-complex (Figure 1F, Supplemental Table S1) ^38–40^. A STRING analysis for functional and physical interactions among proteins revealed a tight interaction network among the U1 snRNA-associated proteins (p-value < 1.0e-16, Supplemental Figure S1) ^41^. Enrichment analysis showed that U1 snRNA-associated proteins feature often RNA binding motifs, helicases and WD40-repeats (Supplemental Data Set 2). Although U1 snRNA-associated proteins were mainly involved in splicing, also other biological processes such as RNA transport, RNA silencing or the regulation of transcription or chromatin assembly were significantly enriched among U1 snRNA-associated proteins (Supplemental Data Set 2). Taken together, the U1 IP-MS experiment revealed more than 200 proteins, statically or dynamically associated with the U1 snRNA and our results suggest functions of the plant U1 snRNP beyond splicing.

### The Arabidopsis U1 snRNP is essential for plant development and transcriptome integrity

To study the functions of the Arabidopsis U1 snRNP and its possible function beyond splicing, the research community lacks plants with reduced levels of core U1 proteins, which cause drastic phenotypic alterations. To address this issue, we generated U1 snRNP knockdown lines through the use of artificial microRNAs (amiRNAs) that targeted the two U1 core subunits U1-70K and U1-C specifically (referred to as *amiR-u1-70k* and *amiR-u1-c*, Figure 2A). This resulted in a reduction of their mRNA levels to approximately 10% of that found in WT plants (Figure 2B, C). We speculated that targeting two different genes encoding proteins forming a common complex would result in similar mutant phenotypes. Indeed, the knockdown of the core U1 subunits U1-C and U1-70K resulted in plants exhibiting pleiotropic defects in plant development, including dwarfism and abnormal leaf development (Figure 2 E-F). While these plants produced a reduced amount of seed, their ability to develop viable seeds despite their extreme phenotype makes them a valuable genetic tool for the entire research community. The altered phenotypes were observed in the vast majority of primary transformants, with the knockdown of *U1-C* always leading to slightly more serve phenotypic alterations (Figure 2 E-F).

**Figure 2:**
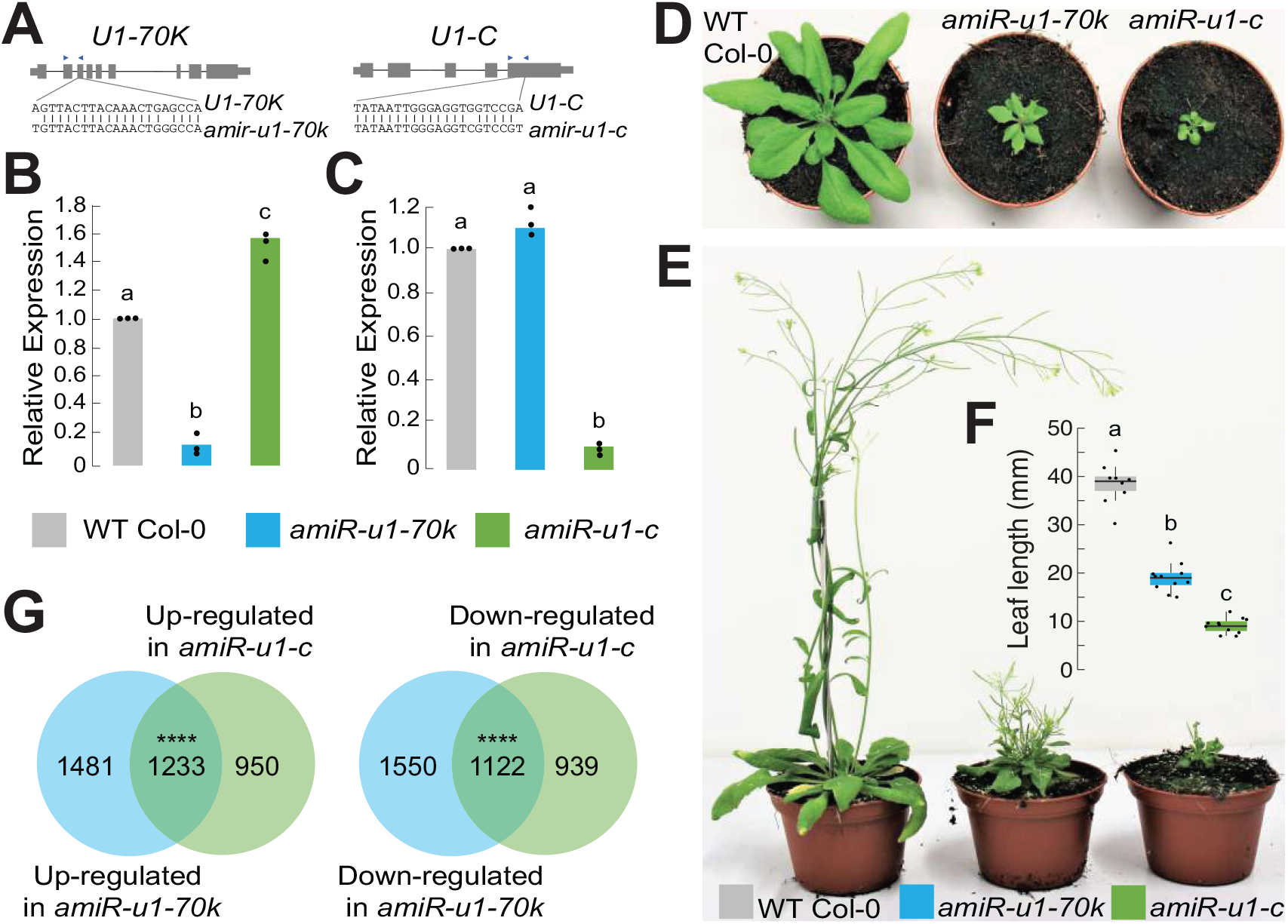
Knock-down of two U1 snRNP core components, U1-70K and U1-C, drastically affects plant development and gene expression. **A:** Gene models of *U1-70K* and *U1-C* and regions used for the design of artificial miRNAs (amiRNAs). The blue rectangle indicates the position of PCR primers used for qPCR in Figure 2B. **B, C:** qRT-PCR analysis of *U1-70K* and *U1-C* levels in seven-day-old WT, *amiR-u1-70k* and *amiR-u1-c* seedlings. The bars indicate the average relative expression in three biological replicates, dots represent indicate the three independent measurements. The letters indicate the statistical significance tested using ANOVA followed by Tukey’s honestly significant difference test (Tukey’s HSD) for pairwise comparison with a significance threshold of p < 0.05. **D, E:** Gross phenotypes of WT, *amiR-u1-70k,* and *amiR-u1-c* plants grown for 21 days (D) or 56 days (E) under long day (16h light/8h darkness) conditions. **F:** Leaf length of WT, *amiR-u1-70k,* and *amiR-u1-c* plants, measured after 21 days. The bars indicate the average leaf length, dots represent indicate individual leaf length measurements. The letters indicate the statistical significance tested using ANOVA followed by Tukey’s honestly significant difference test (Tukey’s HSD) for pairwise comparision with a significance threshold of p < 0.05. **G:** Venn diagrams depicting the overlap of differentially expressed genes in *amiR-u1-70k* and *amiR-u1-c* compared to WT. Expression was determined by RNA-seq and differentially expressed were considered all genes that significantly differed between WT and U1 knockdown line (p_adjusted_ < 0.05).

To determine whether the reduction of *U1-70K* and *U1-C* expression also had comparable effects on the transcriptome, we performed a short-read RNA-seq experiment using WT, *amiR-u1-70k*, and *amiR-u1-c* plants with two to three replicate measurements. In total, we found 2,714 and 2,183 significantly up-regulated and 2,672 and 2,061 significantly down-regulated genes *amiR-u1-70k* and *amiR-u1-c* lines, respectively, when compared to WT plants (Supplemental Data Set 3). A significant number of up-(1233) and down-regulated (1122) genes overlap between *amiR-u1-70k* and *amiR-u1-c* plants (Figure 2G), which further supports the idea that knocking down two different genes encoding proteins of the U1 snRNP result in similar molecular phenotypes.

Because U1-70K and U1-C likely fulfill key functions during splicing, we globally evaluated splicing changes in *amiR-u1-70k* and *amiR-u1-c* lines using the above-described short-read RNA-seq data set and rMATS for bioinformatics analysis ^42^. Alternative splicing events were grouped into different categories: exon skipping, alternative 5’SS or 3’SS, intron retention and mutually exclusive exons (Figure 3A). The largest number of affected transcripts belonged to the exon skipping category (Figure 3A,B, Supplemental Data Set 4). U1 knockdowns in metazoans or mutants in U1 accessory factors such as LUC7 show very similar patterns in splicing defects ^19,24,43^, which is likely due to the altered connection between the U1 and U2 snRNP. More than half of the significant exon skipping events detected were shared between the *amiR-u1-70k* and *amiR-u1-c* lines, which again strongly suggests that both independent knockdown lines have highly similar defects (Figure 3A,B, Supplemental Data Set 4). Also, other splicing defects were found in the U1 knockdown lines which shows that an intact U1 is essential for splicing fidelity in general (Figure 3A,B, Supplemental Data Set 4). The changes in alternative splicing were not due to the mRNA abundance and expression, because we found no significant overlap between alternatively spliced mRNA and differential gene expression (Supplemental Data Set 5). To confirm some of the splicing defects detected using rMATS, we performed RT-PCR with different biological replicates and primers flanking regions of alternative splicing events found in both U1 knockdown lines (Figure 3C). In addition, we performed Oxford Nanopore Technologies (ONT) direct RNA-seq with additional biological replicates of WT, *amiR-u1-70k,* and *amiR-u1-c* plants. While the total number of reads obtained by direct RNA-seq was too low to perform global splicing analysis, the coverage plots of selected splicing events clearly confirmed the short-read RNA-seq analysis (Figure 3D).

**Figure 3:**
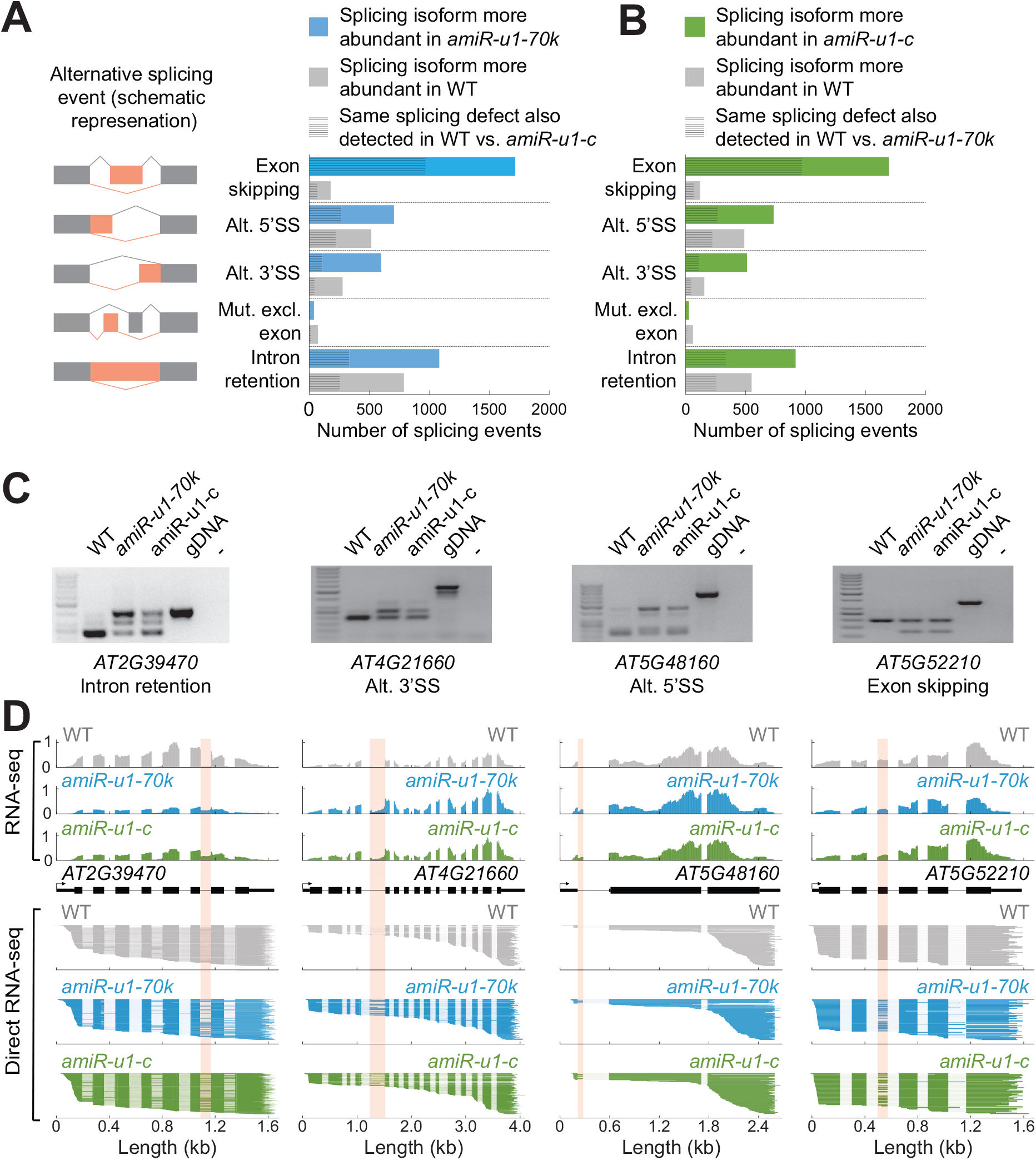
Knock-down of U1-*70K* or *U1-C* causes overlapping splicing defects. **A, B:** Changes in the splicing pattern were calculated based on RNA-seq data from WT, *amiR-u1-70k,* and *amiR-u1-c* plants using rMATS. Splicing changes were subcategorized into exon skipping, alternative 5’splice site (alt. 5’SS), alternative 3’splice site (alt. 3’SS), mutually exclusive exons (mut. Excl. exon), and intron retention. A schematic representation of the different splicing changes is shown in A. **C:** RT-PCR analysis of selected alternative splicing events detected in the RNA-seq data set. Primers used for amplification were designed to flank the splicing event. **D:** ONT direct RNA-seq reads aligned to the genes that produced alternative spliced RNAs (C). The coverage plot of one representative replicate of the RNA-seq data set used for rMATS analysis (A, B) is shown. Pink boxes indicate the alternative splicing events detected by rMATS.

Taken together, these results show the importance of the U1 snRNP in maintaining the normal development of plants and highlight the significance of the U1 snRNP for transcriptome integrity and splicing fidelity. Furthermore, U1 knockdown lines might serve as a powerful tool for studying functions of the Arabidopsis U1 snRNP beyond splicing.

### The Arabidopsis U1 snRNP associates with components of the cleavage and polyadenylation complex (CPSF)

Our U1 IP-MS implied that the Arabidopsis U1 snRNP fulfills additional functions beyond splicing, which we now could address using the U1 knockdown lines. Among the 214 U1 snRNA-associated proteins we identified by U1 IP-MS, we found several cleavage and polyadenylation factors (CPAFs), including components of the cleavage and polyadenylation complex (CPSF) (Figure 4A, Supplemental Table S1). The CPSF recognizes the polyadenylation signal (in metazoans AAUAAA), cleaves the pre-mRNA, and recruits poly(A)polymerases for polyadenylation ^44^. CPSF acts in concert with other complexes such as the Cleavage Stimulation Factor (CstF), Cleavage Factor I, and Cleavage Factor II (CFI and CFII) ^45^. These complexes bind additional cis-regulatory elements, including upstream sequence elements (USE) and downstream sequence elements (DSE). While the canonical polyadenylation signal motif AAUAAA is less well-conserved in plants, which possess a variety of A and U-rich elements, the proteins involved in cleavage and polyadenylation remain highly conserved ^46–48^. The CPSF consists of several subunits: CPSF73, CPSF160, CPSF30, WDR33, FIP1, and CPSF100. CPSF73 functions as an endonuclease and is encoded by two essential genes in Arabidopsis, CPSF73-I and CPSF73-II ^49–51^. FY is the WDR33 homolog in Arabidopsis and recognizes the PAS in concert with CPSF160 ^52–54^. Mutations in the CPSF components showed mild to drastic phenotypic alterations and changes in mRNA cleavage and polyadenylation ^49–51,55–60^.

**Figure 4:**
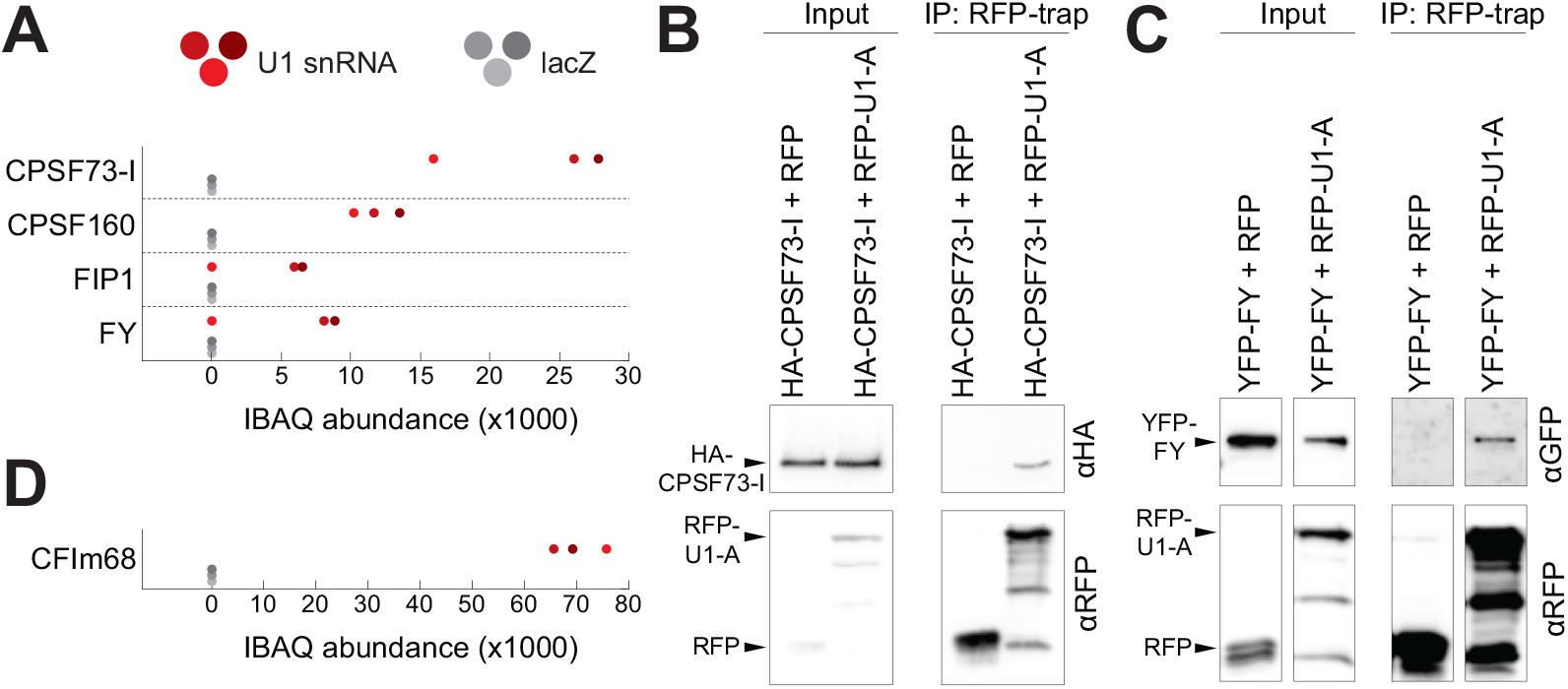
The U1 snRNP associated with components of mRNA cleavage and polyadenylation complexes. **A, D:** Abundance of cleavage and polyadenylation factors (CPAFs) in U1-IP-MS experiments. The three red and grey dots represent intensity-based absolute quantification (IBAQ) values of three biological replicates using the U1 or the lacZ antisense oligonucleotide, respectively. **B, C:** U1-A translationally fused to RFP was coexpressed with an HA-tagged CFSF73-I (B) or a YFP-tagged FY (C) in *Nicotiana benthamiana* plants for transient protein expression. RFP alone served as a negative control. Proteins were isolated and immunoprecipitated using an RFP-affinity matrix. Input and immunoprecipitated fractions (IP) were subjected to protein blot analysis using RFP, HA, and YFP-specific antibodies. Unprocessed blots are available in Figure S2.

We found CPSF73-I, CPSF160, and FIP1 among the 214 significant proteins identified by U1 snRNA IP-MS, suggesting that U1 snRNP forms a high-order complex with the CPSF (Figure 4A). To check this notion, we tested whether protein components of the U1 snRNP co-immunoprecipitate with the CPSF. For this, we transiently co-expressed RFP-U1-A and HA-CPSF73-I fusion proteins and performed affinity purifications using an anti-RFP affinity matrix. HA-CPSF73-I co-purified with RFP-U1-A, but not RFP, which suggests a physical interaction between proteins of the U1 snRNP and the CPSF (Figure 4B). The U1 snRNA IP-MS also contained peptides for two other CPSF subunits, WDR33/FY and CPSF30, but failed to reach the significance threshold (Figure 4A, Supplemental Data Set 1). Still, we also found that YPF-FY co-immunoprecipitated with RFP-U1-A, but not with RFP (Figure 4C). These co-immunoprecipitations of HA-CPSF73-I with RFP-U1-A and YPF-FY with RFP-U1-A, as well as the presence of CPSF73-I, CPSF160, and FIP1 in the U1-IP-MS experiments, strongly support the physical interaction between the Arabidopsis U1 snRNP and CPSF. The U1-IP-MS also retrieved a component of the CFI complex, AtCFI68, which bind to the USE (Figure 4D). This suggests that the U1 snRNP may interact with other components involved in cleavage and polyadenylation, in addition to CPSF components.

### The Arabidopsis U1 snRNP features telescripting function and promotes the selection of canonical polyadenylation sites at the 3’-ends of genes

Given the association of the Arabidopsis U1 snRNP with CPSF components, we investigated its potential role in regulating transcript cleavage and polyadenylation. To address this, we utilized the above-described U1 knock-down lines and performed 3’-end mRNA sequencing with WT, *amiR-u1-70k,* and *amiR-u1-c* plants. In this data set, we could detect approximately 18,000 genes that undergo alternative polyadenylation (APA), with the majority of genes exhibiting more than 4 polyadenylation sites (Supplemental Data Set 6). Changes in the usage of the polyadenylation site were categorized into enhanced and repressed alternative polyadenylation (APA) events. The term “enhanced APA” refers to cases where proximal polyA site usage is higher in WT than in the U1 knockdown lines (Figure 5A), while the term “repressed APA” indicates that the usage of the proximal polyA site in WT is lower than in the U1 knockdown lines (Figure 5A). We found 467 enhanced and 484 repressed PAS sites in *amiR-u1-70k* plants, and 507 enhanced and 693 repressed PAS sites in *amiR-u1-c* plants (Figure 5B,C, Supplemental Data Set 6). Among these, a significant number of enhanced (176, p-value: 6.71e-67) and repressed (102, p-value: 1.24e-06) PAS were shared between *amiR-u1-c* and *amiR-u1-70k* lines, suggesting that U1-C and U1-70K serve similar functions in PAS utilization (Figure 5B, C).

**Figure 5:**
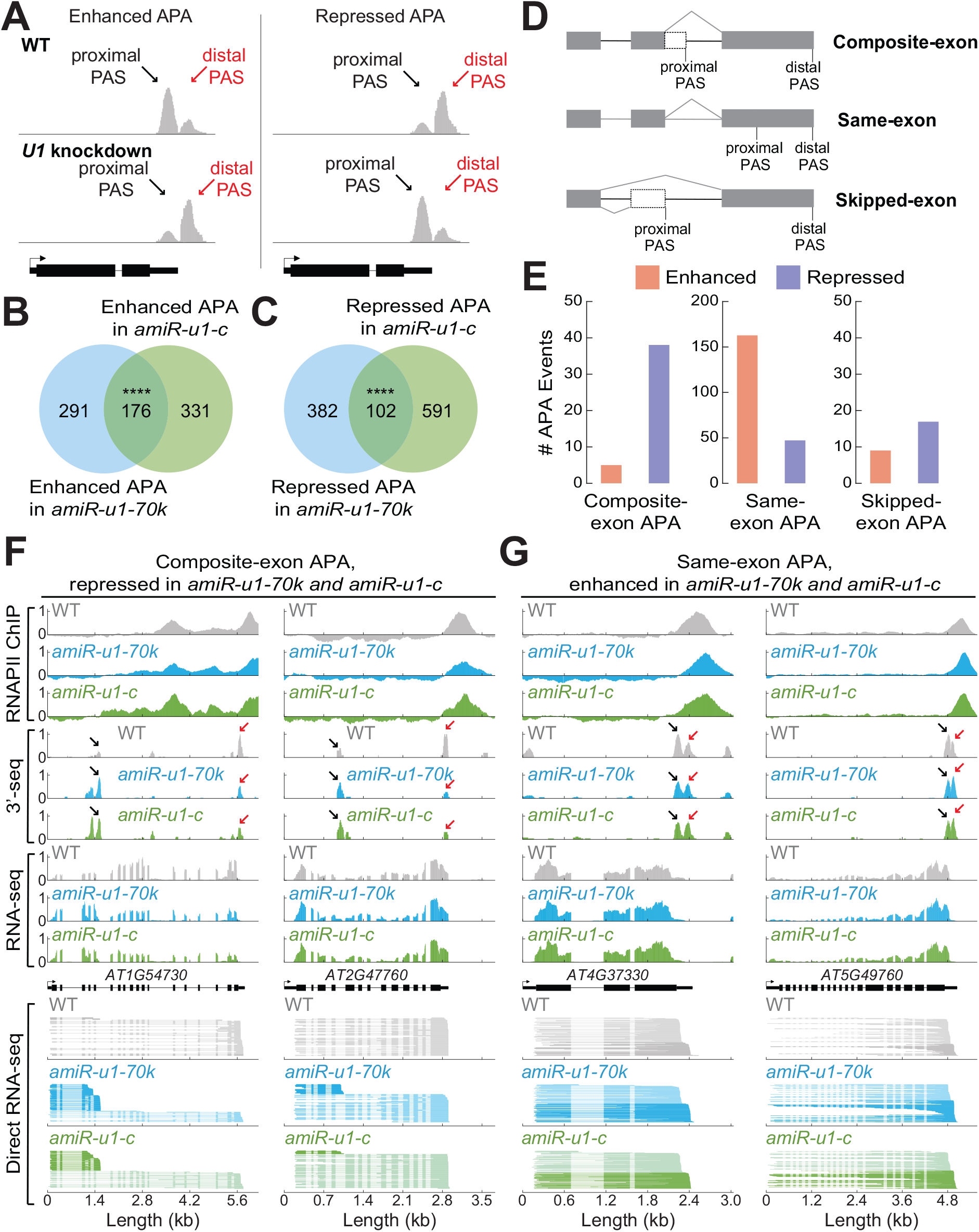
The U1 snRNP regulates alternative polyadenylation in Arabidopsis. **A:** A schematic representation of enhanced and repressed alternative polyadenylation (APA). In enhanced APA events, the proximal PAS site is preferentially utilized. In repressed APA events, the distal PAS site is preferentially utilized. Black arrows indicate the proximal PAS and red arrows indicate the distal PAS. **B, C:** Polyadenylation sites were detected by 3’-end sequencing of RNAs (3’-seq) experiments using RNA isolated from seven-day-old WT, *amiR-u1-70k,* and *amiR-u1-c* seedlings. Venn diagrams depict the overlap of enhanced (B) or repressed (C) APA sites in *amiR-u1-70k* and *amiR-u1-c* when compared to WT. **D:** A schematic representation of three different types of APA, same-exon APA, composite-exon APA, and skipped exon APA. **E:** Number of different APA events detected in both *amiR-u1-70k* and *amiR-u1-c* plants, when compared to WT. Same-exon APA, composite-exon APA, and skipped-exon APA events were further divided into enhanced and repressed events. **F:** Two examples of composite-exon APA events, that are repressed in *amiR-u1-70k* and *amiR-u1-c* plants. The figure depicts the gene models and the corresponding of coverage plots for 3’-seq, RNA-seq, direct RNA-seq, and polymerase II association (RNAPII ChIP). **G:** Two examples of same-exon APA events are enhanced in *amiR-u1-70k* and *amiR-u1-c* plants. The figure depicts the gene models and the corresponding of coverage plots for 3’-seq, RNA-seq, direct RNA-seq, and polymerase II association (RNAPII ChIP).

We further categorized the APA events into three different categories (Figure 5D) ^61^: First, proximal and distal polyA-sites are located in the “same exon”. Second, “composite exon APA” describe APA events that are located on distinct exons. This category should include, e.g. premature polyadenylation events in annotated introns generated by the lack of telescripting. Third, “skipped exon” events refer to alternative polyadenylation events in which the proximal poly(A)site is located in a skipped exon when the distal poly(A) site is utilized. We observed interesting trends for “composite exon” and “same exon” APA events, but no pronounced trend in the *“*skipped-exon” category was found in *amiR-u1-70k* and *amiR-u1-c* plants (Figure 5E, Figure S2 Supplemental Data Set 6).

Both U1 knockdown lines exhibited more repressed than enhanced “composite exon” APA events, indicating that the proximal PAS was utilized more frequently utilized than the distal PAS in U1 knockdown lines (Figure S3, Supplemental Data Set 6). The repressed “composite exon” APA events also significantly overlapped (38 events, p-value: 1.19e-24) between *amiR-u1-70k* (279 events) *and amiR-u1-c* lines (93 events), suggesting that both U1 components target a common set of genes for this type of APA regulation (Figure 5E, Supplemental Data Set 6). Additionally, we detected an accumulation of shorter transcript isoforms for the selected significant repressed “composite exon” APA events by ONT direct RNA-seq (exemplified in Figure 5F). While these shorter transcripts were also detectable in WT plants, they accumulated to higher levels in both U1 knockdown lines (Figure 5F). These results suggest that Arabidopsis genes can generate shorter mRNA through premature polyadenylation in introns, but that the U1 snRNP telescripting function represses the pervasive usage of such premature PAS, akin to the telescripting function of the U1 snRNP in metazoans.

A closer look at the same exon APA events revealed a slightly different picture: Both U1 knockdown lines exhibited more enhanced than repressed “same exon” APA events, and these events significantly overlapped between both knockdown lines (Figure 5E, Supplemental Data Set 6). The increased usage of more distal PAS in this subset of genes led to the generation of longer mRNAs in U1 knock-down lines. These results show that upon U1 knockdown in Arabidopsis, a subset of genes utilized more distal poly(A) sites in terminal exons to produce longer mRNAs (exemplified in Figure 5G). These results suggest at least two distinct functions of the U1 snRNP during polyadenylation: First, the Arabidopsis U1 snRNP suppresses premature polyadenylation in gene bodies through telescripting. Second, the Arabidopsis U1 snRNP promotes the selection of proximal, canonical polyadenylation sites at the 3’-end of mRNAs.

### Usage of distal polyA sites in U1 knockdown might be linked to the release of RNAPII

Two models explain how transcription by RNAPII can be terminated. The allosteric model proposes that transcription of the PAS induces a structural change leading to termination. The torpedo model suggests that after RNA cleavage, the 5’-3’ exonuclease XRN2 rapidly degrades the remaining RNAPII-associated RNA causing termination. More recent data suggest a combined model, in which structural changes facilitate catch-up of RNAPII by XRN2 ^62^. Consistently, the knockdown of factors such as human XRN2 or CPSF73 results in the production of longer transcripts and pile up RNAPII further downstream of the PAS^62,63^.

Since we observed increased usage of distal PAS in terminal exon at some genes upon U1 knockdown, we asked whether RNAPII termination is also affected in U1 knockdown lines. To test this, we performed RNAPII Chromatin Immunoprecipitation (ChIP) experiment experiments followed by sequencing (ChIP-seq) with WT and U1 knockdown lines. At genes with enhanced “same exon” events in *amiR-u1-70k* and *amiR-u1-c*, RNAPII piled up downstream of the RNAPII peak at 3’-ends observed in WT (Figure 6A, exemplified for individual genes in Figure 5G). These results indeed suggest that RNAPII terminates more downstream at this subset of genes upon U1 knockdown. We observed a similar trend for many more genes, although the 3’-end sequencing did not detect any changes in PAS usage between U1 knockdown lines and WT (exemplified in Figure 6B). We therefore decided to investigate the RNAPII distribution among all Arabidopsis genes in WT and U1 knockdowns, irrespectively whether utilizing distal PAS in the same exon accumulates in U1 knockdowns. Also, the RNAPII globally observed a shift of the RNAPII accumulation in *amiR-u1-70k* and *amiR-u1-c* lines. We observed a pronounced RNAPII shift to more distal sites and reduced accumulation of RNAPII at 3’-ends of genes in *amiR-u1-70k* and *amiR-u1-c* lines (Figure 6C), which might suggest a general role of the Arabidopsis U1 snRNP in transcription termination. The reason why we did not detect longer mRNAs when RNAPII terminate more shifts to more distal sites might be the lack of utilizable PAS site or the fact that long 3’UTRs of mRNAs trigger non-sense mediated mRNA decay (NMD) ^64,65^. Then, the full consequences of U1 knockdown on the Arabidopsis transcriptome might only be detectable in U1 knock-down plants, which are also impaired in NMD or other RNA quality control mechanisms.

**Figure 6.**
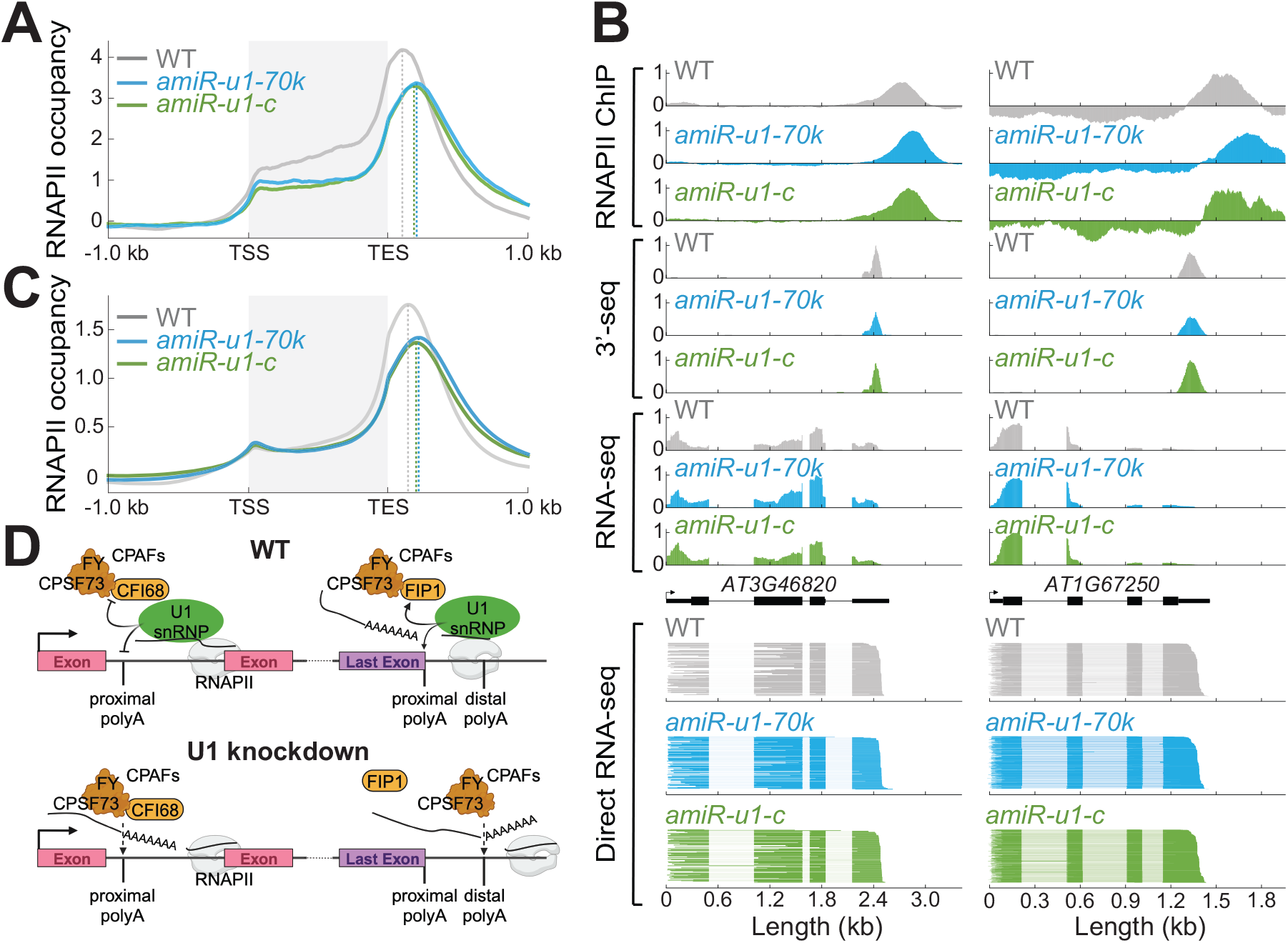
The U1 snRNP affects the distribution of RNA polymerase II at the 3’-end of genes. **A:** Meta plot analysis of RNAPII binding in WT, *amiR-u1-70k,* and *amiR-u1-c* plants to all genes that utilize more distal polyA in 3’UTRs in *amiR-u1-70k* and *amiR-u1-c* plants (same-exon APA events, that are enhanced in *amiR-u1-70k* and *amiR-u1-c* plants, see Figure 5F for individual examples). **B:** Two examples of genes that exhibit a shift of RNAPII accumulation at the 3’-end, but the mRNA of which are not subjected to APA. Along with the polymerase II association (RNAPII ChIP) at each locus, coverage plots for 3’-seq, RNA-seq, and direct RNA-seq polymerase II are shown. **C:** Meta-plot analysis of RNAPII binding to all genes in WT, *amiR-u1-70k,* and *amiR-u1-c* plants. **D:** Proposed model for the function of the U1 snRNP in RNA 3’ processing. The U1 snRNP associates with cleavage and polyadenylation factors (CPAFs), including CPSF77-I, FY, CFI68, and FIP1. These interactions prevent premature intronic polyadenylation or ensure the usage of proximal polyadenylation sites in the last exons. In the absence of U1 snRNP function, intronic polyadenylation occurs, and more distal polyadenylation sites are utilized in the last exons.

## DISCUSSION

In this work, we report the identification of U1 snRNP-associated proteins in Arabidopsis. Using an RNA-centric approach, we enriched known U1 snRNP core and accessory components and identified proteins that may indirectly associate with the U1 snRNP, potentially hinting at their role in mRNA splicing. In general, RNA-centric approaches for the isolation of RNA-containing protein complexes might be powerful tools for the detection mRNPs. For the sake of fairness, one has to admit that the U1 snRNA is a very abundant RNA species, which makes RNA IP-MS experiments for such classes of RNA easier than for lower abundant RNA species. But in general, such RNA-centric approaches to find regulators of RNA processing are attractive, as they do not require generation of transgenics, but might require optimization. For low abundant RNAs, alternative approaches which involve RNA labeling might be better alternatives ^66–70^.

Our *U1-70K* and *U1-C* knockdown lines exhibited much stronger phenotypic alterations compared to previously reported *U1-A* and *U1-70K* T-DNA insertion lines ^20,21^. One possible explanation is that the analyzed T-DNA mutants are not strong or true knock-out alleles, especially for *U1-70K*, where two different T-DNA lines with insertions at the 5’ and 3’-ends of the *U1-70K* gene were studied ^20,21^. Another explanation for the lack of drastically altered phenotypes in U1 T-DNA lines might be functional redundancy. U1-A and U2B’’, both of which bind to the U1 and U2 snRNA stem-loop, respectively, recently evolved from a single ancestral protein and exhibit functional redundancy in metazoans ^71,72^. The sequences of Arabidopsis U1-A and U2B’’ proteins are highly similar ^73^, which might suggest some redundancy also in plants. Although U2B’’ does not bind U1 snRNA under standard conditions ^20,74^, U1 snRNA might be bound by U2B’’ (or other sequence-related U1-A proteins) in *u1-a* mutants in vivo, thus explaining the lack of drastically altered phenotypes in *u1-a* mutants compared to our *U1-70K* and *U1-C* knockdown lines. Antisense morpholino oligonucleotides are a powerful tool to study U1 snRNP functions in human cell culture systems, but similar tools are currently unavailable in the plant research community ^27^. Reduction of *U1-70K* and *U1-C* expression in Arabidopsis by amiRNAs resulted in overlapping phenotypic, RNA expression, splicing, and cleavage/polyadenylation defects. Thus, these amiRNA lines (and further developments using tissue-specific or inducible promoters) become important tools for the future analysis of U1 functions in plants.

The availability of U1-IP-MS data and U1 knockdown lines enabled us to study the function of the Arabidopsis U1 snRNP in mRNA 3’-end processing. Similar to the metazoan U1 snRNP, the Arabidopsis U1 snRNP interacts with RNA 3’-end processing complexes and possesses telescripting function to suppress intronic PAS sites. Moreover, the Arabidopsis U1 snRNP also promotes the usage of proximal PAS sites in 3’-UTRs, which might be linked to a general RNAPII termination defect upon U1 knockdown. The mechanism behind how the Arabidopsis U1 snRNP suppresses intronic PAS while promoting proximal PAS in 3’UTRs remains to be investigated. An RNAi screen in mouse cells shows that the knockdown of CPAFs results in contrasting effects on mRNA length, suggesting that some CPAFs promote while others inhibit certain PAS sites ^75^. For example, the knockdown of AtCFI68 results in shorter mRNAs, while the knockdown of FIP1 increases mRNA length ^75^. In line with this, CFIm68 has been shown as an activator of premature polyadenylation and it was proposed that the U1 snRNP might prevent CFIm68 association to proximal PAS ^36^. Interestingly, we found that AtCFI68 and FIP1 associate with the Arabidopsis U1 snRNP, which might suggest that several U1-CPAF complexes with distinct activities exist in Arabidopsis. Depending on the composition of these complexes and the position along the gene, U1 snRNP suppresses CPAF activities, while a U1 snRNP with distinct protein partner enhances cleavage and polyadenylation at proximal sites in 3’UTRs (Figure 4D). Identification of cis and trans factors responsible for the distinct modes of U1 action will be an interesting subject for future studies.

Alternative polyadenylation plays a pivotal role in gene expression control in plants and several factors involved in APA have been described in plants ^52,53,76–80^. The U1 snRNP has not yet been linked to APA in plants and while the changes we analyzed occur only in an artificial condition (U1 knockdown), we think that the U1 telescripting function and U1 3’UTR length regulation are also important layers of adaptive gene regulation in plants. Early reports suggested that cleavage and polyadenylation within introns is rare ^81,82^, but usage of intronic PAS to regulate gene expression in Arabidopsis has been described in several instances; such alternative polyadenylation might lead to non-functional RNAs to control the abundance of the canonical mRNA or to generation alternative mRNAs encoding alternative protein ^83–86^. Modulation of U1 snRNP telescripting function to regulate APA might therefore add an important layer for gene expression in Arabidopsis and in crops, too ^87,88^. Whether certain condition globally affects telescripting in plants, such as in humans under heat-shock conditions, remains to be elucidated ^34^.

## MATERIALS AND METHODS

### Plant material and growth conditions

All *Arabidopsis thaliana* lines used in this study were in the Columbia (Col-0) background. Plants on soil were grown under a long-day cycle (16 hours light/8 hours night) at 22°C/20°C conditions. For seedlings grown on plates, seeds were first surface sterilized with 80% ethanol containing 0.05% Triton X-100. Afterward, seeds were grown on half-strength Murashige Skoog (MS) plates containing 0.8% phytoagar and grown for 7 days (for RNA-sequencing) or 14 days (for ChIP or ChIRP) under continuous light conditions at 22°C.

For the construction of artificial microRNA against *U1-70K* and *U1-C*, oligonucleotides (Supplemental Table 3) were derived from (http://wmd3.weigelworld.org/cgi-bin/webapp.cgi?page=Home;project=stdwmd). PCR products were amplified using Phusion High Fidelity DNA Polymerase (NEB) and pRS300 plasmid containing the miR319a precursor served as the template ^89,90^. The engineered artificial microRNAs were subcloned into pCR8/GW/TOPO (Thermo Scientific) and transferred into a Gateway Cloning system pGWB602 ^91^ using Gateway LR Clonase II Enzyme Mix (Thermo Scientific). The resulting plasmids were transformed into the Agrobacterium strain GV3101 and introduced into Arabidopsis Col-0 plants through the floral dipping method ^92^.

### RNA extractions, RT-qPCR and Illumina library preparation

Total RNA was extracted using Direct-zol^TM^ RNA Miniprep (Zymo Research) according to the manufacturer’s instructions. For the validation of the alternative splicing defects, 1-2 µg of RNA was treated with DNAse I (ThermoFisher Scientific), and the cDNA was prepared using the RevertAid First Strand cDNA Synthesis kit (ThermoFisher Scientific) using 100 µM oligodT. RT-PCR was performed using the Dreamtaq DNA Polymerase (ThermoFisher Scientific) and run on 2% agarose gel. For the RT-qPCR experiments, we used Maxima SYBR Green (ThermoFischer Scientific) in Bio-Rad CFX-384 and calculated the relative expression using the 2(-ΔΔCT) with PP2A as the control. All the oligonucleotides are listed in Supplementary Table 3. For the RNA-sequencing experiments, 5 µg of RNA was treated with DNAse I (ThermoFisher Scientific) and cleaned up using the RNA Clean and Concentrator^TM^-5 (Zymo Research). PolyA mRNA was isolated using the NEBNext Poly(A) mRNA Magnetic Isolation Module (New England Biolabs). Afterward, the cDNA libraries were prepared using the NEBNext Ultra Directional RNA Library Prep Kit for Illumina (New England Biolabs). The resulting libraries were measured using the Qubit dsDNA High Sensitivity Assay Kit (Thermo Fisher Scientific) and size distribution was determined using the Agilent High Sensitivity DNA Kit (Agilent). Libraries were pooled together for paired-end sequencing on an Illumina Hi-Seq 3000. For 3’-end RNA sequencing, DNAse-treated RNA was sent to Lexogen GmbH (Vienna, Austria) for library construction using the Quantseq 3’ mRNA-seq Library Prep kit REV.

### Differential gene expression and alternative splicing analysis

Paired end reads were trimmed using Trim Galore (version 0.6.7) (https://github.com/FelixKrueger/TrimGalore) with Cutadapt ^93^ (version 3.4) and filtered by aligning all reads to the tRNA and rRNA transcripts of *Arabidopsis thaliana*. For that purpose, the latest transcriptome (ATRTD3) was queried for tRNA and rRNA transcripts using the functional descriptions provided by Araport11 ^94^. The reads were then aligned to the tRNA/rRNA reference using HISAT2 (version 2.2.1) ^95^. Reads that did not align to tRNA and rRNA were used for downstream analysis. Quality control was performed before and after trimming and filtering with fastQC (version 0.11.9) (https://www.bioinformatics.babraham.ac.uk/projects/fastqc/) and multiQC (version 1.13) ^96^. Filtered and trimmed reads were quantified at transcript level with salmon (version 1.9.0) using ATRTD3 ^94,97^. Quantified transcript level reads were summarized to gene level and imported to R (version 4.2.2) using tximport (version 1.26.1) ^98^). Differentially expressed genes (p < 0.05) were called using the R package DESeq2 (version 1.38.3) ^99^. Additional packages used for the analysis and visualization are ggrepel (version 0.9.3, https://github.com/slowkow/ggrepel), ggplot2 (version 3.4.2) ^100^ and dplyr (version 1.1.2) (https://github.com/tidyverse/dplyr). For a full session report and additional quality control plots refer to the jupyter notebook provided within the Github repository https://github.com/WeberJoachim/Mangilet_et_al_2023. For the analysis of differentially spliced transcripts the filtered and trimmed reads were mapped to the Arabidopsis genome (version TAIR10)^101^ with HISAT2 (version 2.2.1). The resulting alignments were converted to BAM format, sorted and indexed using SAMtools (version 1.9)^102^. Differentially spliced transcripts were identified from indexed and sorted BAM files with rMATS (version 4.1.2). An additional software used in this analysis is seqkit (version 2.3.1) ^103^. All workflows and specific parameters used in this analysis are available under https://github.com/WeberJoachim/Mangilet_et_al_2023. Pipelines were implemented using Nextflow ^104^, programs were used as Singularity containers ^105^. Singularity image files were pulled from Galaxy ^106^.

### 3’-end mRNA sequencing analysis

3’-end mRNA sequencing reads were analyzed using the apa toolkit within the expressRNA framework ^61^. For visualization, the reads were trimmed using Trim Galore and filtered by aligning to a rRNA / tRNA reference using HISAT2. The filtered and trimmed reads were aligned to TAIR10 using HISAT2, resulting alignment files were converted to BAM format, sorted and indexed using SAMtools. BAM files were then converted to BedGraph using deepTools and merged using UCSC WiggleTools (version 1.2.8) ^107^. Merged BedGraph files were further used for visualization.

### Nanopore direct RNA-sequencing

Total RNA was isolated using RNAzol® RT (Sigma-Aldrich, R4533) from three biological replicates of WT, *amiR-u1-70k* and *amiR-u1-c* seedlings. according to the manufactures instructions and quantified using NanoDrop® ND-1000 Spectrophotometer. We isolated polyA RNA using the Ambion™ Poly(A)Purist™ MAG K kit (Thermo Fisher Scientific, AM1922) according to manufactures instructions. Quantity and quality of total and polyA-selected RNA were determined using the Qubit RNA HS assay and 2100 Agilent Bioanalyzer using the Agilent RNA 6000 Pico kit. For direct RNA-seq library preparation, the SQK-RNA002 kit (Oxford Nanopore Technologies) was used together with NEBNext® Quick Ligation Reaction Buffer (NEB B6058), T4 DNA Ligase 2M U/ml (NEB M0202), SuperScript III Reverse Transcriptase (Thermo Fisher Scientific, 18080044) and Agencourt RNAClean XP beads according to manufactures instructions. Qubit 1x dsDNA HS assay was used to quantitate 1 µl of the library, and the remainder was loaded on a primed PromethION flow cell (FLO-PRO002 R9) and run on a PromethION. Fast5 files were basecalled using Cuda (version 12.1.0) and Guppy (Version 6.2.1), with the statistical model “rna_r9.4.1_70bps_hac_prom.cfg”. Initial quality analysis was performed using FastQC and summarized with multiQC. Basecalled reads were aligned against the genome (TAIR10) using minimap2 (version 2.24)^108^. SAM files were converted to BAM, sorted and indexed using SAMtools (version 1.17). Because of variation of library sizes ranging from 0.1 to 2.6 million reads within replicates, the alignments from all three biological replicates were merged using SAMtools to perform qualitative analysis depicted in Figure 5E and 6B.

### Comprehensive Identification of RNA-binding proteins (ChIRP)

The original protocol was adapted from ^37^ with some minor modifications. Nine grams of 14-day-old Arabidopsis Col-0 seedlings were harvested and crosslinked with 3% formaldehyde for 15 minutes under a vacuum chamber at 85 kPa. Vacuum infiltration was repeated once more to ensure proper cross-linking. The cross-linking reaction was then quenched by adding 4 mL of 1.25 M Glycine for 5 minutes in the vacuum. Cross-linked seedlings were then washed three times with distilled water, dried on blotting paper, and stored at −80°C. To isolate the nuclei, frozen materials were grounded with liquid Nitrogen and resuspended in HONDA buffer (400 mM Sucrose, 1.25% Ficoll, 2.5% Dextran, 25 mM HEPES-KOH pH 7.4, 10 mM MgCl_2_, 0.5% Triton X-100, 1 mM PMSF, cOmplete Protease Inhibitor Cocktail EDTA-free [Roche], and 10 mM DTT). The homogenate was passed through two layers of Miracloth and was centrifuged at 1500 g for 15 minutes at 4°C. The pellet was then carefully washed five times with HONDA buffer until most of the green material was removed at 1500g for 5 minutes at 4°C centrifuge. Then washed again with M3 buffer (10 mM Sodium phosphate pH 7.0, 100 mM NaCl, 10 mM DTT, and 1X Protein Inhibitor). Finally, the pellet was resuspended in sonic buffer (10 mM Sodium Phosphate pH7.0, 100 mM Sodium chloride, 0.5% Sarkosyl, 10 mM EDTA, 1X Complete cocktail, 1 mM PEFA). Chromatin shearing was done using the Covaris S220 under the following conditions: 20% Duty Cycle, 140 Peak intensity, 200 Cycles per burst, and a total of 3 minutes of cycle time. The samples were centrifuged at 13000 rpm for 5 minutes at 4°C. The supernatant was then transferred into a DNA LoBind tube (Eppendorf), flash frozen in liquid nitrogen and stored at −80°C. Chromatin was thawed at room temperature together with the probes for U1 snRNA and control (Supplemental Table 3). Fifty microliters of chromatin served as the protein input. Two ml of hybridization buffer (750 mM NaCl, 50 mM Tris-HCl pH 7.0, 1mM EDTA, 1% SDS, 15% Formamide, 1x Protease Inhibitor, 1x PMSF, 1x Riboblock (40 U/µl) [ThermoFisher Scientific], Plant Specific Protease Inhibitor [Sigma]). One microliter of the 100 µM probe was added and allowed to gently rotate end-to-end at 37°C for 4 hours in a hybridization oven. With 2 hours remaining for the hybridization, 100 µl of Dynabeads MyOne Streptavidin C1 (ThermoFisher Scientific) were prepared by removing the storage buffer and washing them three times with 1 mL of unsupplemented nuclear lysis (50 mM Tris-HCl, 10 mM EDTA, 1% SDS) buffer using a magnetic stand.. When the hybridization was finished, 100 µl of the washed beads were added to the mixture and incubated for an additional 30 minutes. During this incubation, the wash buffer (2x SSC, 0.5 % SDS) was prepared and pre-warmed at 37°C before use. When the bead binding is completed, the mixture was briefly centrifuged and the beads were separated from the mixture for two minutes in a magnetic stand. One microliter of the Wash Buffer was used to wash the beads and gently rotated again at 37°C for 5 minutes in a hybridization oven. The washing step is repeated four times, for a total of five washes. For the last wash, all the buffer was removed. For the preparation for the mass spectrometry analysis, the beads were washed three times in 20 mM sodium bicarbonate buffer.

### Protein on beads digestion

All steps for protein digestion were performed at room temperature as described before ^109^. Briefly, beads were resuspended in denaturation buffer (6 M urea, 2 M thiourea, 10 mM Tris buffer, pH 8.0), and proteins were reduced and subsequently alkylated by incubation in 1 mM dithiothreitol (DTT) for one hour followed by addition of 5.5mM iodacetamide (IAA) for another hour in the dark. Proteins were pre-digested with LysC for three hours at pH 8. Beads were then diluted in 4 volumes 20 mM ammonium bicarbonate buffer and proteins digested with 2 µg trypsin per estimated 100 µg protein at pH 8 overnight. Acidified peptides were desalted with C18 stage tips as described previously ^110^.

### Mass spectrometry

LC-MS/MS analyses of eluted samples were performed on an Easy nano-LC (Thermo Scientific) coupled to an LTQ Orbitrap XL mass spectrometer (Thermo Scientific) as decribed elsewhere ^111^. The peptide mixtures were injected onto the column in HPLC solvent A (0.1% formic acid) at a flow rate of 500 nl/min and subsequently eluted with a 49 minute segmented gradient of 10–33-50-90 % of HPLC solvent B (80% acetonitrile in 0.1% formic acid) at a flow rate of 200 nl/min. The 15 most intense precursor ions were sequentially fragmented in each scan cycle using collision-induced dissociation (CID). In all measurements, sequenced precursor masses were excluded from further selection for 30 s. The target values were 5000 charges for MS/MS fragmentation and 10^6^ charges for the MS scan. Due to high contamination of polymers in the samples it was decided to further purify the samples via PHOENIX Peptide Clean-up Kit (PreOmics) according to user manual. Final measurements were performed after PHOENIX Kit purification as described above.

### Mass spectrometry data processing

The MS data of all runs together were processed with MaxQuant software suite v.1.5.2.8 ^112^. A database search was performed using the Andromeda search engine which, is integrated into MaxQuant ^113^. MS/MS spectra were searched against a target-decoy Uniprot database from *A. thaliana* downloaded 2019-02-13 consisting of 91,457 protein entries from *A. thaliana* and 245 commonly observed contaminants. In a database search, full specificity was required for trypsin. Up to two missed cleavages were allowed. Carbamidomethylation of cysteine was set as a fixed modification, whereas oxidation of methionine and acetylation of protein N-terminus were set as variable modifications. Initial mass tolerance was set to 4.5 parts per million (ppm) for precursor ions and 0.5 daltons (Da) for fragment ions. Peptide, protein, and modification site identifications were reported at a false discovery rate (FDR) of 0.01, estimated by the target/decoy approach ^114^. Match between runs was enabled for samples within one group so for U1, U2, and control samples separately. iBAQ and LFQ settings were enabled. MaxQuant data were analyzed using msVolcano for the detection of significantly enriched proteins using the following parameters ^115^: FDR = 0.04; curvature = 0.75; min. fold change = 0 or FDR = 0.05; curvature = 2.5; min. fold change = 0 for U1-IP-MS and U2-IP-MS, respectively.

### Co-Immunoprecipitation

For the expression of HA-, RFP- or YFP-tagged proteins, the coding sequence of each protein was PCR-amplified and subcloned into pCR™8/GW/TOPO® (Invitrogen). To generate binary plasmids, the entry vectors were recombined using Gateway™ LR Clonase™ II (Thermo Scientific) with either pGWB642 for the expression of YFP-tagged fusion proteins, pGWB515 for the expression of HA-tagged fusion proteins or pGWB654 for the expression of RFP fusion proteins ^91^. Binary plasmids were transformed into *Agrobacterium tumefaciens* (strain GV3101). Proteins were expressed by Agrobacterium-mediated transient expression in *Nicotiana benthamiana*. For this, Agrobacterium was grown overnight at 28°C and cultures were pelleted by centrifugation. The pellet were resuspended in infiltration media (10 mM MgCl_2_, 10 mM MES-KOH, pH 5.6 and 100 µM acetosyringone) and the OD_600_ was adjusted to 0.5. After incubated for three hours at 22°C with light agitation. one or two leaves per *N. benthamiana* plant were infiltrated with infiltration mixture. After three days, transformed tobacco leaves were snap-frozen, grounded to a fine powder and resuspended in the protein lysis buffer (50 mM Tris HCl pH7.4, 150 mM NaCl, 10% Glycerol, 0.5 % TritonX-100, 0.5 % Nonidet^TM^ P 40 Substitute, 1 mM PMSF, 2 mM DTT, 50 µM MG132, Plant specific protease inhibitor (Sigma-Aldrich P9599), and cOmplete Protease Inhibitor Cocktail EDTA-free (Roche). After centrifugation at 13000 x g for 10 minutes at 4°C. the supernatant was used for immunoprecipitation. For each immunoprecipitation, 20 µl of RFP-trap beads (Chromotek) were equilibrated by washing them three times with wash buffer (50 mM Tris HCl pH 7.5, 150 mM NaCl, 10 % Glycerol). The protein samples were added to the equilibrated beads and incubated for 1 hour on a rotating wheel at 4°C. The input samples were incubated together with the IP samples. After incubation, the beads were washed three times with wash buffer before incubation in Laemmli buffer at 80°C for 10 minutes. The isolated proteins were resolved by SDS-PAGE, blotted to nitrocellulose membranes and incubated with antibodies specific for GFP (Chromotek, 3h9), RFP (Chromotek, 6g6) or HA (Agrisera; AS12 2200). An HRP-conjugated secondary antibody (rat AS10 1115, rabbit AS09 602 and mouse AS10 1115, all Agrisera) and the Western Bright Chemiluminescence Substrate Sirius (Biozym) was used for protein detection.

### Chromatin Immunoprecipitation (ChIP)

The method is adapted from ^116^. Three grams of 14-day old Arabidopsis seedlings were collected and fixed with 40 mL 1% formaldehyde in MQ buffer (10 mM Sodium phosphate pH7.0, 50 mM Sodium chloride) for 10 minutes under a vacuum chamber at 85 kPa. Vacuum infiltration was repeated once more to ensure proper cross-linking. The cross-linking reaction was then quenched by adding 4 mL of 1.25M Glycine for 5 minutes in the vacuum. Cross-linked seedlings were then washed three times with distilled water, dried on paper, and stored at −80C. To isolate the nuclei, frozen materials were then grounded with liquid Nitrogen and resuspend in HONDA buffer (400 mM Sucrose, 1.25% Ficoll, 2.5% Dextran, 25 mM HEPES-KOH pH7.4, 10 mM MgCl2, 0.5% Triton X-100, 1 mM PMSF, Proteinase Inhibitor cocktail, and 10 mM DTT). The resuspended plant materials were filtered with 2 layers of Miracloth and transferred into a new 50 mL tube. The homogenate was centrifuged at 1500g for 15 minutes at 4C. The pellet was then carefully washed five times with HONDA buffer until most of the green material was removed at 1500 g for 5 minutes at 4C centrifuge. Then washed again with M3 buffer (10 mM Sodium phosphate pH7.0, 100mM Sodium chloride, 10 mM DTT and 1X Protein Inhibitor). Then the pellet was resuspended in Sonic buffer (10 mM Sodium Phosphate pH7.0, 100 mM Sodium chloride, 0.5% Sarkosyl, 10 mM EDTA, 1X Complete cocktail, 1 mM PEFA). Chromatin shearing was done using the Covaris S220 under the following conditions: 20% Duty Cycle, 140 Peak intensity, 200 Cycles per burst, and a total of 3 minutes of cycle time. The samples were then centrifuged at 13000rpm for 5 minutes at 4C. The supernatant was then transferred into a DNA lobind tube.

For the immunoprecipitation experiment, 700 µl of the solubilized chromatin was used, and 140 µl for the input. And then IP buffer (50 mM HEPES pH7.4, 150 mM KCl, 5 mM MgCl_2_, 0.01 mM ZnSO_4_, 1% Triton X-100, 0.05% SDS) was added to the IP and input. RNAPII CTD (Abcam, ab817) were added to the IP and incubated overnight in a rotating wheel overnight at 4°C. The following day, 40 µl of Protein A/G agarose beads (Santa Cruz Biotechnology, Cat. No. sc2001) was added to the IP and incubated for 6 hours in a rotating wheel at 4°C. After the incubation, beads were pelleted by centrifugation and washed five times with 1mL of IP buffer on a rotating wheel with centrifugation in each wash. Associated DNA with the proteins was then eluted with 120 µl of cold acidic glycine buffer pH 2.8 (100 mM Glycine, 500 mM NaCl, 0.05 % Tween-20, HCl). The supernatant was transferred to a tube containing 150 µl of Tris pH 9.0. This elution with glycine was repeated twice and each elution was transferred into the same tube. RNAse A was added and incubated at 37°C for 15 minutes. To denature the proteins, 1.5 µl of Proteinase K was added and incubated overnight at 37C. A second aliquot of Proteinase K was added to the samples and incubated at 65C for 6 hours to reverse the crosslinking. DNA was then purified using MinElute (Qiagen) according to the manufacturer’s instructions with minor modifications. The IP samples were then divided into 2 and 3 volumes of ERC buffer was added. The pH was adjusted using 3 M sodium acetate. The mixture was then added to the spin column and washed with the PE buffer and eluted with 35 µl EB buffer. ChIP DNA libraries were prepared using the NEBNext Ultra II DNA Library Prep Kit for Illumina (New England Biolabs) according to the manufacturer’s instructions. The libraries were prepared without size selection. Multiplexing was done using the NEBNext Multiplex Oligos for Illumina (Set 1, 2,3,4). The concentration of the libraries was determined using the Qubit TM dsDNA HS Assay Kit (Thermo Fisher Scientific) and size distribution was measured using the Agilent High Sensitivity DNA Kit (Agilent). Libraries were pooled together and performed paired-end sequencing on an Illumina Hi-Seq 2000.

### ChIP-seq analysis

Paired end reads from ChIP-seq (Chromatin Immuno Precipitation DNA-Sequencing) were trimmed using Trim Galore. Trimmed reads were then aligned to the Arabidopsis genome (version TAIR10) ^101^ with HISAT2 using the “--no-splice-alignment” option. Mapped reads were further analyzed with the MACS2 (version 2.9.1.)^117^. Therefore, the IGG control pileups was subtracted from the treatment and input control pileups. The resulting pileups (BedGraphs) were then compared using fold enrichment between IGG corrected treatment and input. Quality control of pileups was performed by converting BedGraphs to bigWig files and subsequent multibigwigsummary and plotCorrelation using deepTools (version 3.5.2)^118^. Here it was discovered that replicate 1 of *amiR-u1-c* behaves differently from all other samples and thus was discarded in the downstream analysis. Meta plots were assembled by merging the bigWig files and then plotting them using deepTools plotProfile.

### Data visualization

For visualizing all sequencing reads, we created a fork of the long-read visualization framework from FLEP-seq ^119^ and added the functionality to plot BedGraph files. The code can be found in the Jupyter notebook within the Github repo of this study or as standalone repository under https://github.com/WeberJoachim/Viz_bdg_and_nanopore_bam.

### Data availability

All raw data sets, along with metadata files, are publicly available at ENA or PRIDE under the accession numbers PRJEB65251 (for RNA and DNA sequencing) or PXD045484 (for proteomic analyses). All analysis pipelines and parameters applied are accessible at https://github.com/WeberJoachim/Mangilet_et_al_2023.

## Supporting information

Supplemental_Data_Set_1

Supplemental_Data_Set_2

Supplemental_Data_Set_3

Supplemental_Data_Set_4

Supplemental_Data_Set_5

Supplemental_Data_Set_6

## ACKNOWLEDGEMENTS

We are grateful to Udo Gowik for his help in the initial phase of this project. We greatly appreciate the valuable comments and critical reading of the manuscript by Cornelius Schmidtke. This work was founded by the German Science Foundation (DFG), grant LA2633-4/2, to S.L..

## AUTHOR CONTRIBUTIONS

A.F.M and S.L. designed the study. A.F.M, S.Sc., I.D.-B. performed experiments. J.W., G.R. A.F.M, I.D.-B., B.M. and S.L. analyzed the data, S.St. and T.S. contributed analytical tools, A.F.M and S.L. wrote the article with contributions from all authors.

## REFERENCES

1 Will, C. L. & Luhrmann, R. Spliceosome structure and function. Cold Spring Harb Perspect Biol 3, doi:10.1101/cshperspect.a003707 (2011).

2 Chen, W. & Moore, M. J. Spliceosomes. Curr Biol 25, R181–183, doi:10.1016/j.cub.2014.11.059 (2015).

3 Kondo, Y., Oubridge, C., van Roon, A. M. & Nagai, K. Crystal structure of human U1 snRNP, a small nuclear ribonucleoprotein particle, reveals the mechanism of 5’ splice site recognition. Elife 4, doi:10.7554/eLife.04986 (2015).

4 Li, Q. Q., Liu, Z., Lu, W. & Liu, M. Interplay between Alternative Splicing and Alternative Polyadenylation Defines the Expression Outcome of the Plant Unique OXIDATIVE TOLERANT-6 Gene. Sci Rep 7, 2052, doi:10.1038/s41598-017-02215-z (2017).

5 Plaschka, C., Lin, P. C., Charenton, C. & Nagai, K. Prespliceosome structure provides insights into spliceosome assembly and regulation. Nature 559, 419–422, doi:10.1038/s41586-018-0323-8 (2018).

6 Guiro, J. & O’Reilly, D. Insights into the U1 small nuclear ribonucleoprotein complex superfamily. Wiley Interdiscip Rev RNA 6, 79–92, doi:10.1002/wrna.1257 (2015).

7 Lerner, M. R. & Steitz, J. A. Antibodies to small nuclear RNAs complexed with proteins are produced by patients with systemic lupus erythematosus. Proc Natl Acad Sci U S A 76, 5495–5499, doi:10.1073/pnas.76.11.5495 (1979).

8 Zhang, D. & Rosbash, M. Identification of eight proteins that cross-link to pre-mRNA in the yeast commitment complex. Genes Dev 13, 581–592, doi:10.1101/gad.13.5.581 (1999).

9 Puig, O., Gottschalk, A., Fabrizio, P. & Seraphin, B. Interaction of the U1 snRNP with nonconserved intronic sequences affects 5’ splice site selection. Genes Dev 13, 569–580, doi:10.1101/gad.13.5.569 (1999).

10 Forch, P., Puig, O., Martinez, C., Seraphin, B. & Valcarcel, J. The splicing regulator TIA-1 interacts with U1-C to promote U1 snRNP recruitment to 5’ splice sites. Embo J 21, 6882–6892, doi:10.1093/emboj/cdf668 (2002).

11 Espinosa, S. et al. Human PRPF39 is an alternative splicing factor recruiting U1 snRNP to weak 5’ splice sites. RNA 29, 97–110, doi:10.1261/rna.079320.122 (2022).

12 Wang, L. et al. The RNA-binding protein RBP45D of Arabidopsis promotes transgene silencing and flowering time. Plant J 109, 1397–1415, doi:10.1111/tpj.15637 (2022).

13 Chang, P., Hsieh, H. Y. & Tu, S. L. The U1 snRNP component RBP45d regulates temperature-responsive flowering in Arabidopsis. Plant Cell 34, 834–851, doi:10.1093/plcell/koab273 (2022).

14 Stepien, A. et al. Chromatin-associated microprocessor assembly is regulated by the U1 snRNP auxiliary protein PRP40. Plant Cell 34, 4920–4935, doi:10.1093/plcell/koac278 (2022).

15 Hernando, C. E. et al. A Role for Pre-mRNA-PROCESSING PROTEIN 40C in the Control of Growth, Development, and Stress Tolerance in Arabidopsis thaliana. Front Plant Sci 10, 1019, doi:10.3389/fpls.2019.01019 (2019).

16 Wang, C. et al. The Arabidopsis thaliana AT PRP39-1 gene, encoding a tetratricopeptide repeat protein with similarity to the yeast pre-mRNA processing protein PRP39, affects flowering time. Plant Cell Rep 26, 1357–1366, doi:10.1007/s00299-007-0336-5 (2007).

17 Kanno, T. et al. A Genetic Screen for Pre-mRNA Splicing Mutants of Arabidopsis thaliana Identifies Putative U1 snRNP Components RBM25 and PRP39a. Genetics 207, 1347–1359, doi:10.1534/genetics.117.300149 (2017).

18 Huang, W. et al. A genetic screen in Arabidopsis reveals the identical roles for RBP45d and PRP39a in 5’ cryptic splice site selection. Front Plant Sci 13, 1086506, doi:10.3389/fpls.2022.1086506 (2022).

19 de Francisco Amorim, M., et al. The U1 snRNP Subunit LUC7 Modulates Plant Development and Stress Responses via Regulation of Alternative Splicing. Plant Cell 30, 2838–2854, doi:10.1105/tpc.18.00244 (2018).

20 Gu, J. et al. Spliceosomal protein U1A is involved in alternative splicing and salt stress tolerance in Arabidopsis thaliana. Nucleic Acids Res 46, 1777–1792, doi:10.1093/nar/gkx1229 (2018).

21 Chen, M. X. et al. Phylogenetic comparison of 5’ splice site determination in central spliceosomal proteins of the U1-70K gene family, in response to developmental cues and stress conditions. Plant J 103, 357–378, doi:10.1111/tpj.14735 (2020).

22 Golovkin, M. & Reddy, A. S. Expression of U1 small nuclear ribonucleoprotein 70K antisense transcript using APETALA3 promoter suppresses the development of sepals and petals. Plant Physiol 132, 1884–1891, doi:10.1104/pp.103.023192 (2003).

23 Salz, H. K. et al. The Drosophila U1-70K protein is required for viability, but its arginine-rich domain is dispensable. Genetics 168, 2059–2065, doi:10.1534/genetics.104.032532 (2004).

24 Rösel, T. D. et al. RNA-Seq analysis in mutant zebrafish reveals role of U1C protein in alternative splicing regulation. Embo J 30, 1965–1976, doi:10.1038/emboj.2011.106 (2011).

25 Baserga, S. J. & Steitz, J. A. The Diverse World of Small Ribonucleoproteins. Cold Spring Harbor Monograph Series (1993).

26 Almada, A. E., Wu, X., Kriz, A. J., Burge, C. B. & Sharp, P. A. Promoter directionality is controlled by U1 snRNP and polyadenylation signals. Nature 499, 360–363, doi:10.1038/nature12349 (2013).

27 Kaida, D. et al. U1 snRNP protects pre-mRNAs from premature cleavage and polyadenylation. Nature 468, 664–668, doi:10.1038/nature09479 (2010).

28 Berg, M. G. et al. U1 snRNP determines mRNA length and regulates isoform expression. Cell 150, 53–64, doi:10.1016/j.cell.2012.05.029 (2012).

29 Mimoso, C. A. & Adelman, K. U1 snRNP increases RNA Pol II elongation rate to enable synthesis of long genes. Mol Cell 83, 1264–1279 e1210, doi:10.1016/j.molcel.2023.03.002 (2023).

30 Yin, Y. et al. U1 snRNP regulates chromatin retention of noncoding RNAs. Nature 580, 147–150, doi:10.1038/s41586-020-2105-3 (2020).

31 Venters, C. C., Oh, J. M., Di, C., So, B. R. & Dreyfuss, G. U1 snRNP Telescripting: Suppression of Premature Transcription Termination in Introns as a New Layer of Gene Regulation. Cold Spring Harb Perspect Biol 11, doi:10.1101/cshperspect.a032235 (2019).

32 Oh, J. M. et al. U1 snRNP telescripting regulates a size-function-stratified human genome. Nat Struct Mol Biol 24, 993–999, doi:10.1038/nsmb.3473 (2017).

33 Waldrop, M. A. et al. Intron mutations and early transcription termination in Duchenne and Becker muscular dystrophy. Hum Mutat 43, 511–528, doi:10.1002/humu.24343 (2022).

34 Cugusi, S. et al. Heat shock induces premature transcript termination and reconfigures the human transcriptome. Mol Cell 82, 1573–1588 e1510, doi:10.1016/j.molcel.2022.01.007 (2022).

35 Oh, J. M. et al. U1 snRNP regulates cancer cell migration and invasion in vitro. Nat Commun 11, 1, doi:10.1038/s41467-019-13993-7 (2020).

36 So, B. R. et al. A Complex of U1 snRNP with Cleavage and Polyadenylation Factors Controls Telescripting, Regulating mRNA Transcription in Human Cells. Mol Cell 79, doi:10.1016/j.molcel.2019.08.007 (2019).

37 Chu, C. et al. Systematic discovery of Xist RNA binding proteins. Cell 161, 404–416, doi:10.1016/j.cell.2015.03.025 (2015).

38 Raczynska, K. D. et al. The SERRATE protein is involved in alternative splicing in Arabidopsis thaliana. Nucleic Acids Res 42, 1224–1244, doi:10.1093/nar/gkt894 (2014).

39 Jia, T. et al. The Arabidopsis MOS4-Associated Complex Promotes MicroRNA Biogenesis and Precursor Messenger RNA Splicing. Plant Cell 29, 2626–2643, doi:10.1105/tpc.17.00370 (2017).

40 Laubinger, S. et al. Dual roles of the nuclear cap-binding complex and SERRATE in pre-mRNA splicing and microRNA processing in Arabidopsis thaliana. Proc Natl Acad Sci U S A 105, 8795–8800, doi:10.1073/pnas.0802493105 (2008).

41 Szklarczyk, D. et al. The STRING database in 2023: protein-protein association networks and functional enrichment analyses for any sequenced genome of interest. Nucleic Acids Res 51, D638–D646, doi:10.1093/nar/gkac1000 (2023).

42 Shen, S. et al. rMATS: robust and flexible detection of differential alternative splicing from replicate RNA-Seq data. Proc Natl Acad Sci U S A 111, E5593–5601, doi:10.1073/pnas.1419161111 (2014).

43 Morcos, P. A. Achieving targeted and quantifiable alteration of mRNA splicing with Morpholino oligos. Biochem Biophys Res Commun 358, 521–527, doi:10.1016/j.bbrc.2007.04.172 (2007).

44 Proudfoot, N. J. Ending the message: poly(A) signals then and now. Genes Dev 25, 1770–1782, doi:10.1101/gad.17268411 (2011).

45 Boreikaite, V. & Passmore, L. A. 3’-End Processing of Eukaryotic mRNA: Machinery, Regulation, and Impact on Gene Expression. Annu Rev Biochem 92, 199–225, doi:10.1146/annurev-biochem-052521-012445 (2023).

46 Loke, J. C. et al. Compilation of mRNA polyadenylation signals in Arabidopsis revealed a new signal element and potential secondary structures. Plant Physiol 138, 1457–1468, doi:10.1104/pp.105.060541 (2005).

47 Hunt, A. G., Xing, D. & Li, Q. Q. Plant polyadenylation factors: conservation and variety in the polyadenylation complex in plants. BMC Genomics 13, 641, doi:10.1186/1471-2164-13-641 (2012).

48 Zhang, S. et al. New insights into Arabidopsis transcriptome complexity revealed by direct sequencing of native RNAs. Nucleic Acids Res 48, 7700–7711, doi:10.1093/nar/gkaa588 (2020).

49 Xu, R. et al. The 73 kD subunit of the cleavage and polyadenylation specificity factor (CPSF) complex affects reproductive development in Arabidopsis. Plant Mol Biol 61, 799–815, doi:10.1007/s11103-006-0051-6 (2006).

50 Xu, R., Ye, X. & Quinn Li, Q. AtCPSF73-II gene encoding an Arabidopsis homolog of CPSF 73 kDa subunit is critical for early embryo development. Gene 324, 35–45, doi:10.1016/j.gene.2003.09.025 (2004).

51 Liu, Y. et al. snRNA 3’ End Processing by a CPSF73-Containing Complex Essential for Development in Arabidopsis. PLoS Biol 14, e1002571, doi:10.1371/journal.pbio.1002571 (2016).

52 Simpson, G. G., Dijkwel, P. P., Quesada, V., Henderson, I. & Dean, C. FY Is an RNA 3′ End-Processing Factor that Interacts with FCA to Control the Arabidopsis Floral Transition. Cell 113, 777–787, doi:10.1016/s0092-8674(03)00425-2 (2003).

53 Yu, Z., Lin, J. & Li, Q. Q. Transcriptome Analyses of FY Mutants Reveal Its Role in mRNA Alternative Polyadenylation. Plant Cell 31, 2332–2352, doi:10.1105/tpc.18.00545 (2019).

54 Schonemann, L. et al. Reconstitution of CPSF active in polyadenylation: recognition of the polyadenylation signal by WDR33. Genes Dev 28, 2381–2393, doi:10.1101/gad.250985.114 (2014).

55 Hou, Y. et al. CPSF30-L-mediated recognition of mRNA m(6)A modification controls alternative polyadenylation of nitrate signaling-related gene transcripts in Arabidopsis. Mol Plant 14, 688–699, doi:10.1016/j.molp.2021.01.013 (2021).

56 Song, P. et al. Arabidopsis N(6)-methyladenosine reader CPSF30-L recognizes FUE signals to control polyadenylation site choice in liquid-like nuclear bodies. Mol Plant 14, 571–587, doi:10.1016/j.molp.2021.01.014 (2021).

57 Hong, L. et al. Alternative polyadenylation is involved in auxin-based plant growth and development. Plant J 93, 246–258, doi:10.1111/tpj.13771 (2018).

58 Tellez-Robledo, B. et al. The polyadenylation factor FIP1 is important for plant development and root responses to abiotic stresses. Plant J 99, 1203–1219, doi:10.1111/tpj.14416 (2019).

59 Wang, C. et al. FIP1 Plays an Important Role in Nitrate Signaling and Regulates CIPK8 and CIPK23 Expression in Arabidopsis. Front Plant Sci 9, 593, doi:10.3389/fpls.2018.00593 (2018).

60 Lin, J., Xu, R., Wu, X., Shen, Y. & Li, Q. Q. Role of cleavage and polyadenylation specificity factor 100: anchoring poly(A) sites and modulating transcription termination. Plant J 91, 829–839, doi:10.1111/tpj.13611 (2017).

61 Rot, G. et al. High-Resolution RNA Maps Suggest Common Principles of Splicing and Polyadenylation Regulation by TDP-43. Cell Rep 19, 1056–1067, doi:10.1016/j.celrep.2017.04.028 (2017).

62 Eaton, J. D., Francis, L., Davidson, L. & West, S. A unified allosteric/torpedo mechanism for transcriptional termination on human protein-coding genes. Genes Dev 34, 132–145, doi:10.1101/gad.332833.119 (2020).

63 Fong, N. et al. Effects of Transcription Elongation Rate and Xrn2 Exonuclease Activity on RNA Polymerase II Termination Suggest Widespread Kinetic Competition. Mol Cell 60, 256–267, doi:10.1016/j.molcel.2015.09.026 (2015).

64 Hogg, J. R. & Goff, S. P. Upf1 senses 3’UTR length to potentiate mRNA decay. Cell 143, 379–389, doi:10.1016/j.cell.2010.10.005 (2010).

65 Kertesz, S. et al. Both introns and long 3’-UTRs operate as cis-acting elements to trigger nonsense-mediated decay in plants. Nucleic Acids Res 34, 6147–6157, doi:10.1093/nar/gkl737 (2006).

66 Yi, W. et al. CRISPR-assisted detection of RNA-protein interactions in living cells. Nat Methods 17, 685–688, doi:10.1038/s41592-020-0866-0 (2020).

67 Yang, X. et al. Proximity labeling: an emerging tool for probing in planta molecular interactions. Plant Commun 2, 100137, doi:10.1016/j.xplc.2020.100137 (2021).

68 Qin, W., Cho, K. F., Cavanagh, P. E. & Ting, A. Y. Deciphering molecular interactions by proximity labeling. Nat Methods 18, 133–143, doi:10.1038/s41592-020-01010-5 (2021).

69 Grawe, C., Stelloo, S., van Hout, F. A. H. & Vermeulen, M. RNA-Centric Methods: Toward the Interactome of Specific RNA Transcripts. Trends Biotechnol 39, 890–900, doi:10.1016/j.tibtech.2020.11.011 (2021).

70 Burjoski, V. & Reddy, A. S. N. The Landscape of RNA-Protein Interactions in Plants: Approaches and Current Status. Int J Mol Sci 22, doi:10.3390/ijms22062845 (2021).

71 Saldi, T., Wilusz, C., MacMorris, M. & Blumenthal, T. Functional redundancy of worm spliceosomal proteins U1A and U2B’’. Proc Natl Acad Sci U S A 104, 9753–9757, doi:10.1073/pnas.0701720104 (2007).

72 Delaney, K. J., Williams, S. G., Lawler, M. & Hall, K. B. Climbing the vertebrate branch of U1A/U2B’’ protein evolution. RNA 20, 1035–1045, doi:10.1261/rna.044255.114 (2014).

73 Simpson, G. G. et al. Molecular characterization of the spliceosomal proteins U1A and U2B" from higher plants. Embo J 14, 4540–4550, doi:10.1002/j.1460-2075.1995.tb00133.x (1995).

74 Terzi, L. C. & Simpson, G. G. Arabidopsis RNA immunoprecipitation. Plant J 59, 163–168, doi:10.1111/j.1365-313X.2009.03859.x (2009).

75 Li, W. et al. Systematic profiling of poly(A)+ transcripts modulated by core 3’ end processing and splicing factors reveals regulatory rules of alternative cleavage and polyadenylation. PLoS Genet 11, e1005166, doi:10.1371/journal.pgen.1005166 (2015).

76 Liu, F., Marquardt, S., Lister, C., Swiezewski, S. & Dean, C. Targeted 3’ processing of antisense transcripts triggers Arabidopsis FLC chromatin silencing. Science 327, 94–97, doi:10.1126/science.1180278 (2010).

77 Hornyik, C., Terzi, L. C. & Simpson, G. G. The spen family protein FPA controls alternative cleavage and polyadenylation of RNA. Dev Cell 18, 203–213, doi:10.1016/j.devcel.2009.12.009 (2010).

78 Zhang, Y. et al. Integrative genome-wide analysis reveals HLP1, a novel RNA-binding protein, regulates plant flowering by targeting alternative polyadenylation. Cell Res 25, 864–876, doi:10.1038/cr.2015.77 (2015).

79 Lin, J. et al. HDA6-dependent histone deacetylation regulates mRNA polyadenylation in Arabidopsis. Genome Res 30, 1407–1417, doi:10.1101/gr.255232.119 (2020).

80 Zhang, X. et al. CFI 25 Subunit of Cleavage Factor I is Important for Maintaining the Diversity of 3’ UTR Lengths in Arabidopsis thaliana (L.) Heynh. Plant Cell Physiol 63, 369–383, doi:10.1093/pcp/pcac002 (2022).

81 Wu, X. et al. Genome-wide landscape of polyadenylation in Arabidopsis provides evidence for extensive alternative polyadenylation. Proc Natl Acad Sci U S A 108, 12533–12538, doi:10.1073/pnas.1019732108 (2011).

82 Sherstnev, A. et al. Direct sequencing of Arabidopsis thaliana RNA reveals patterns of cleavage and polyadenylation. Nat Struct Mol Biol 19, 845–852, doi:10.1038/nsmb.2345 (2012).

83 Parker, M. T. et al. Widespread premature transcription termination of Arabidopsis thaliana NLR genes by the spen protein FPA. Elife 10, doi:10.7554/eLife.65537 (2021).

84 Cyrek, M. et al. Seed Dormancy in Arabidopsis Is Controlled by Alternative Polyadenylation of DOG1. Plant Physiol 170, 947–955, doi:10.1104/pp.15.01483 (2016).

85 Duc, C., Sherstnev, A., Cole, C., Barton, G. J. & Simpson, G. G. Transcription termination and chimeric RNA formation controlled by Arabidopsis thaliana FPA. PLoS Genet 9, e1003867, doi:10.1371/journal.pgen.1003867 (2013).

86 Guo, C., Spinelli, M., Liu, M., Li, Q. Q. & Liang, C. A Genome-wide Study of "Non-3UTR" Polyadenylation Sites in Arabidopsis thaliana. Sci Rep 6, 28060, doi:10.1038/srep28060 (2016).

87 Fu, H. et al. Genome-wide dynamics of alternative polyadenylation in rice. Genome Res 26, 1753–1760, doi:10.1101/gr.210757.116 (2016).

88 Zhou, Q. et al. Differential alternative polyadenylation contributes to the developmental divergence between two rice subspecies, japonica and indica. Plant J 98, 260–276, doi:10.1111/tpj.14209 (2019).

89 Ossowski, S., Schwab, R. & Weigel, D. Gene silencing in plants using artificial microRNAs and other small RNAs. Plant J 53, 674–690, doi:10.1111/j.1365-313X.2007.03328.x (2008).

90 Schwab, R., Ossowski, S., Riester, M., Warthmann, N. & Weigel, D. Highly specific gene silencing by artificial microRNAs in Arabidopsis. Plant Cell 18, 1121–1133, doi:10.1105/tpc.105.039834 (2006).

91 Nakagawa, T. et al. Improved Gateway binary vectors: high-performance vectors for creation of fusion constructs in transgenic analysis of plants. Biosci Biotechnol Biochem 71, 2095–2100, doi:10.1271/bbb.70216 (2007).

92 Clough, S. J. & Bent, A. F. Floral dip: a simplified method for Agrobacterium-mediated transformation of Arabidopsis thaliana. Plant J 16, 735–743, doi:10.1046/j.1365-313x.1998.00343.x (1998).

93 Martin, M. Cutadapt removes adapter sequences from high-throughput sequencing reads. EMBnet.journal 17, doi:10.14806/ej.17.1.200 (2011).

94 Zhang, R. et al. A high-resolution single-molecule sequencing-based Arabidopsis transcriptome using novel methods of Iso-seq analysis. Genome Biol 23, 149, doi:10.1186/s13059-022-02711-0 (2022).

95 Kim, D., Paggi, J. M., Park, C., Bennett, C. & Salzberg, S. L. Graph-based genome alignment and genotyping with HISAT2 and HISAT-genotype. Nat Biotechnol 37, 907–915, doi:10.1038/s41587-019-0201-4 (2019).

96 Ewels, P., Magnusson, M., Lundin, S. & Kaller, M. MultiQC: summarize analysis results for multiple tools and samples in a single report. Bioinformatics 32, 3047–3048, doi:10.1093/bioinformatics/btw354 (2016).

97 Srivastava, A., Malik, L., Smith, T., Sudbery, I. & Patro, R. Alevin efficiently estimates accurate gene abundances from dscRNA-seq data. Genome Biol 20, 65, doi:10.1186/s13059-019-1670-y (2019).

98 Soneson, C., Love, M. I. & Robinson, M. D. Differential analyses for RNA-seq: transcript-level estimates improve gene-level inferences. F1000Res 4, 1521, doi:10.12688/f1000research.7563.2 (2015).

99 Love, M. I., Huber, W. & Anders, S. Moderated estimation of fold change and dispersion for RNA-seq data with DESeq2. Genome Biol 15, 550, doi:10.1186/s13059-014-0550-8 (2014).

100. ggplot2. (2016).

101 Lamesch, P. et al. The Arabidopsis Information Resource (TAIR): improved gene annotation and new tools. Nucleic Acids Res 40, D1202–1210, doi:10.1093/nar/gkr1090 (2012).

102 Li, H. et al. The Sequence Alignment/Map format and SAMtools. Bioinformatics 25, 2078–2079, doi:10.1093/bioinformatics/btp352 (2009).

103 Shen, W., Le, S., Li, Y. & Hu, F. SeqKit: A Cross-Platform and Ultrafast Toolkit for FASTA/Q File Manipulation. PLoS One 11, e0163962, doi:10.1371/journal.pone.0163962 (2016).

104 Di Tommaso, P. et al. Nextflow enables reproducible computational workflows. Nat Biotechnol 35, 316–319, doi:10.1038/nbt.3820 (2017).

105 Kurtzer, G. M., Sochat, V. & Bauer, M. W. Singularity: Scientific containers for mobility of compute. PLoS One 12, e0177459, doi:10.1371/journal.pone.0177459 (2017).

106 Galaxy, C. The Galaxy platform for accessible, reproducible and collaborative biomedical analyses: 2022 update. Nucleic Acids Res 50, W345–W351, doi:10.1093/nar/gkac247 (2022).

107 Zerbino, D. R., Johnson, N., Juettemann, T., Wilder, S. P. & Flicek, P. WiggleTools: parallel processing of large collections of genome-wide datasets for visualization and statistical analysis. Bioinformatics 30, 1008-1009, doi:10.1093/bioinformatics/btt737 (2014).

108 Li, H. Minimap2: pairwise alignment for nucleotide sequences. Bioinformatics 34, 3094–3100, doi:10.1093/bioinformatics/bty191 (2018).

109 Mukherjee, J. et al. beta-Actin mRNA interactome mapping by proximity biotinylation. Proc Natl Acad Sci U S A 116, 12863–12872, doi:10.1073/pnas.1820737116 (2019).

110 Rappsilber, J., Mann, M. & Ishihama, Y. Protocol for micro-purification, enrichment, pre-fractionation and storage of peptides for proteomics using StageTips. Nat Protoc 2, 1896–1906, doi:10.1038/nprot.2007.261 (2007).

111 Franz-Wachtel, M. et al. Global detection of protein kinase D-dependent phosphorylation events in nocodazole-treated human cells. Mol Cell Proteomics 11, 160–170, doi:10.1074/mcp.M111.016014 (2012).

112 Cox, J. & Mann, M. MaxQuant enables high peptide identification rates, individualized p.p.b.-range mass accuracies and proteome-wide protein quantification. Nat Biotechnol 26, 1367–1372, doi:10.1038/nbt.1511 (2008).

113 Cox, J. et al. Andromeda: a peptide search engine integrated into the MaxQuant environment. J Proteome Res 10, 1794–1805, doi:10.1021/pr101065j (2011).

114 Elias, J. E. & Gygi, S. P. Target-decoy search strategy for increased confidence in large-scale protein identifications by mass spectrometry. Nat Methods 4, 207–214, doi:10.1038/nmeth1019 (2007).

115 Singh, S., Hein, M. Y. & Stewart, A. F. msVolcano: A flexible web application for visualizing quantitative proteomics data. Proteomics 16, 2491–2494, doi:10.1002/pmic.201600167 (2016).

116 Speth, C. et al. Arabidopsis RNA processing factor SERRATE regulates the transcription of intronless genes. Elife 7, doi:10.7554/eLife.37078 (2018).

117 Zhang, Y. et al. Model-based analysis of ChIP-Seq (MACS). Genome Biol 9, R137, doi:10.1186/gb-2008-9-9-r137 (2008).

118 Ramirez, F. et al. deepTools2: a next generation web server for deep-sequencing data analysis. Nucleic Acids Res 44, W160–165, doi:10.1093/nar/gkw257 (2016).

119 Long, Y., Jia, J., Mo, W., Jin, X. & Zhai, J. FLEP-seq: simultaneous detection of RNA polymerase II position, splicing status, polyadenylation site and poly(A) tail length at genome-wide scale by single-molecule nascent RNA sequencing. Nat Protoc 16, 4355–4381, doi:10.1038/s41596-021-00581-7 (2021).

