## Supplementary material for "The Arabidopsis U1 snRNP regulates mRNA 3’-end processing": Figure_S1

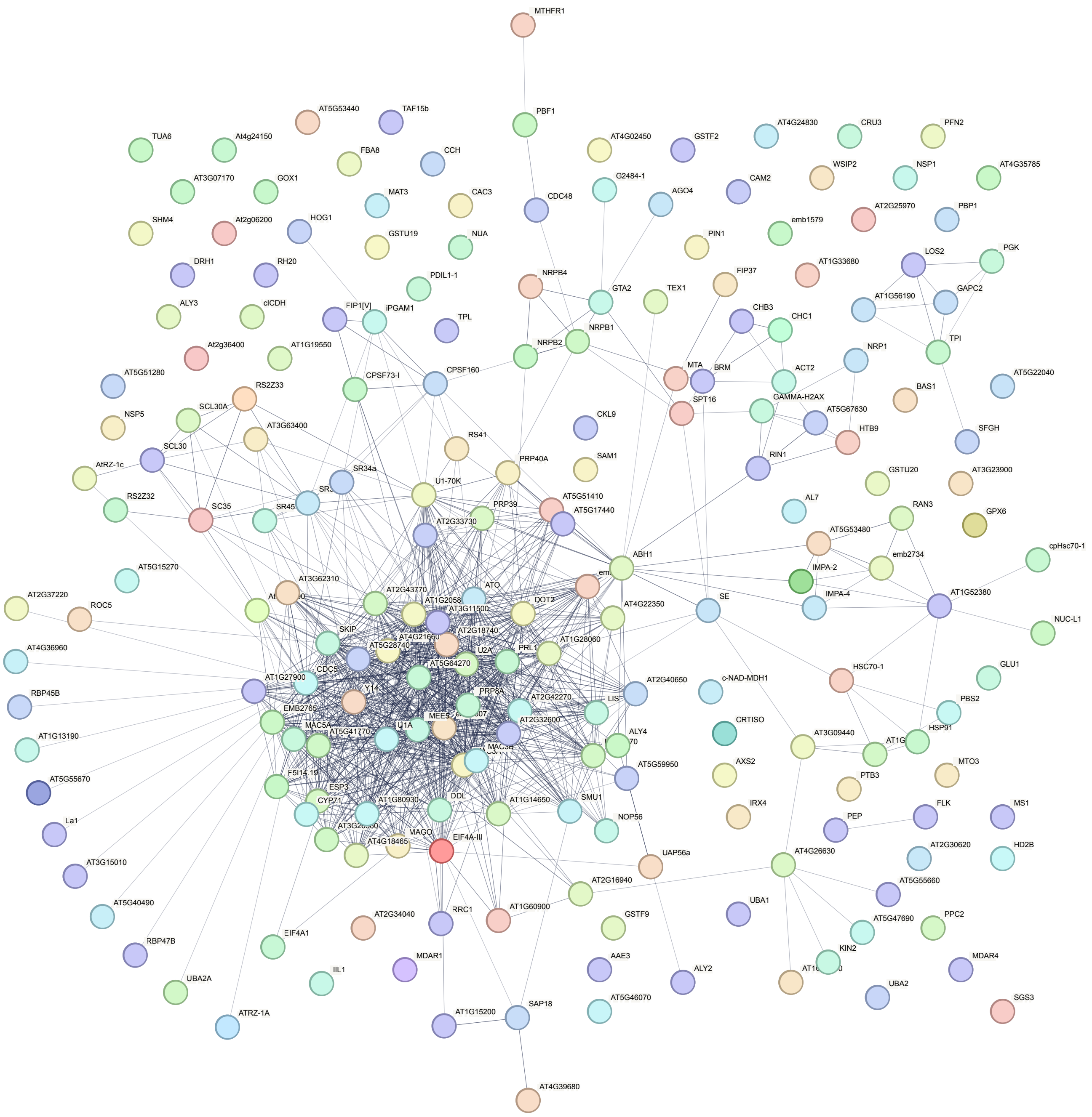


### Figure S1

**Figure S1: String analysis reveals known interactions between significantly enriched proteins in the U1-IP-MS experiment.** We applied the following parameter for the String analysis: interaction sources: Textmining, Experiments, Databases, minimum required interaction score: high confidence.
