## Supplementary material for "The Arabidopsis U1 snRNP regulates mRNA 3’-end processing": Figure_S2

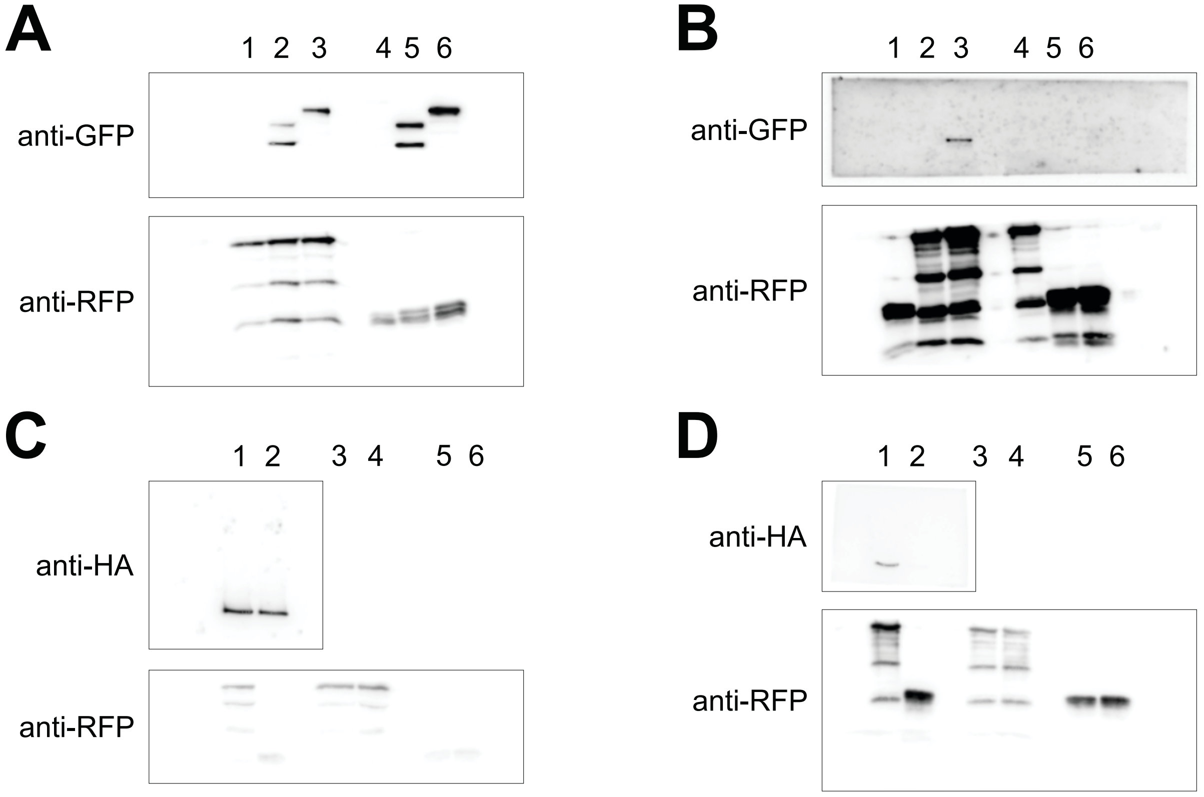


### Figure S2

**Figure S2: Unprocessed blots used for assembly of Figure 4 B and C.**

**A:** Input samples for figure 4B: Lane 3: YFP-FY + RFP-U1A; Lane 6: YFP-FY + RFP.
**B:** IP RFP trap samples for figure 4B: Lane 3: YFP-FY + RFP-U1A; Lane 6: YFP-FY + RFP.
**C:** Input samples for figure 4C. Lane 1: HA-CPSF73-1 + RFP-U1A; Lane 2: HA-CPSF73-1 + RFP
**D:** IP-RFP trap samples for figure 4C. Lane 1: HA-CPSF73-1 + RFP-U1A; Lane 2: HA-CPSF73-1 + RFP
