## Supplementary material for "The Arabidopsis U1 snRNP regulates mRNA 3’-end processing": Figure_S3

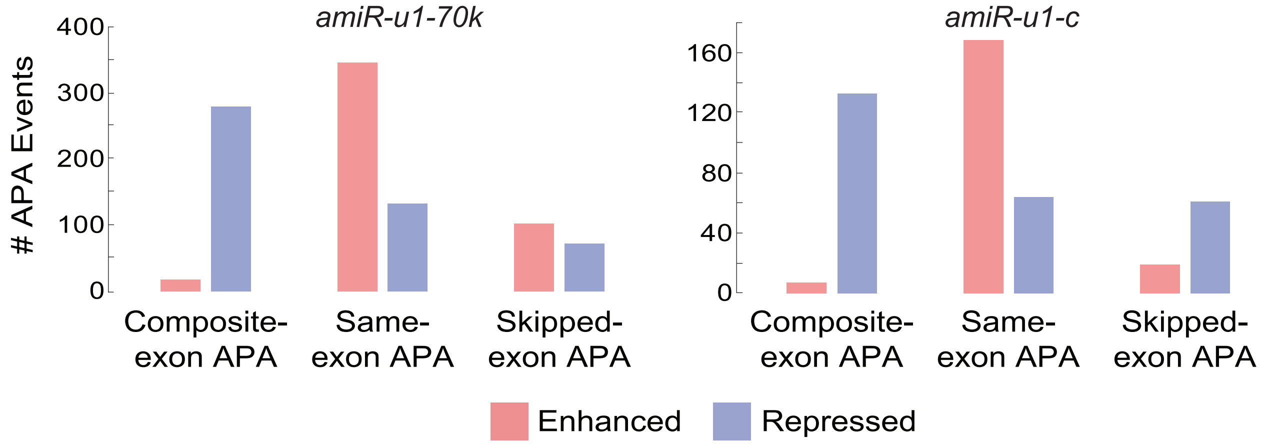


### Figure S3

**Figure S3: APA events in *amiR-u1-70k* and *amiR-u1-c* plants.**
Number of different APA events detected in *amiR-u1-70k* and *amiR-u1-c* plants, when compared to WT. Same-exon APA, composite-exon APA, and skipped-exon APA events were further divided into enhanced and repressed events. The overlap of the different APA events is depicted in Figure 5E.
